# Mapping glycoprotein structure reveals defining events in the evolution of the *Flaviviridae*

**DOI:** 10.1101/2024.02.06.579159

**Authors:** Jonathon C.O. Mifsud, Spyros Lytras, Michael R. Oliver, Kamilla Toon, Vincenzo A. Costa, Edward C. Holmes, Joe Grove

**Affiliations:** Sydney Institute for Infectious Diseases, School of Medical Sciences, The University of Sydney, Sydney, NSW 2006, Australia; MRC-University of Glasgow Centre for Virus Research, Glasgow, G61 1QH, UK; Laboratory of Data Discovery for Health Limited, Hong Kong SAR, China

## Abstract

Viral glycoproteins drive membrane fusion in enveloped viruses and determine host range, tissue tropism and pathogenesis. Despite their importance, there is a fragmentary understanding of glycoproteins within the *Flaviviridae*; for many species the glycoproteins have not yet been identified, for others, such as the hepaciviruses, the molecular mechanisms of membrane fusion remain uncharacterised. Here, we combine comprehensive phylogenetic analyses with systematic protein structure prediction to survey glycoproteins across the entire *Flaviviridae*. We discover class-II fusion systems, homologous to the orthoflavivirus E glycoprotein, in most species, including highly-divergent jingmenviruses and large genome flaviviruses. However, the E1E2 glycoproteins of the hepaci-, pegi- and pestiviruses are structurally distinct, may represent a novel class of fusion mechanism, and are strictly associated with infection of vertebrate hosts. By mapping glycoprotein distribution onto the underlying phylogeny we reveal a complex history of evolutionary events that have shaped the diverse virology and ecology of the *Flaviviridae*.

## Introduction

The *Flaviviridae* is a highly diverse family of enveloped positive-sense (+ve) RNA viruses that includes important pathogens of humans (e.g., Dengue virus [DENV], Zika virus, hepatitis C virus [HCV]) and other animals (e.g., classical swine fever virus, bovine viral diarrhoea virus), as well as many viruses that pose emerging threats to human health (e.g., West Nile virus, Alongshan virus, Haseki tick virus ^1–3^). The *Flaviviridae* is currently classified into four genera: *Orthoflavivirus*, *Pestivirus*, *Pegivirus* and *Hepacivirus* ^4^. In recent years a remarkable diversity of novel flaviviruses with varied genome structures have been discovered, including the jingmenvirus group that are unique in being both segmented and potentially multicomponent ^5,6^. Another group, tentatively known as the large genome flaviviruses (LGF) are primarily associated with invertebrates ^7^, and have also been linked to plants ^8,9^ and vertebrates ^3^. LGFs have genomes up to 39.8kb in length, challenging previous assumptions about the maximum genome size achievable by RNA viruses that lack proofreading mechanisms ^10,11^. These latter two groups have yet to receive taxonomic ratification, and consensus is lacking regarding their placement within flavivirus phylogeny.

Previous efforts to reconstruct the evolutionary history of large and diverse families of RNA viruses such as the *Flaviviridae* have largely relied on the phylogenetic analysis of highly conserved viral proteins, most notably the RNA-dependent RNA polymerase (RdRp)^7,12^. Although of considerable utility, the functions and features that define virus biology and pathogenesis are typically encoded by highly divergent sequences outside of the conserved replication machinery. In these regions it is difficult to detect deep sequence homology and hence perform reliable multiple sequence alignment or phylogenetic analysis. As a consequence, our understanding of long-term virus evolution is generally based on the analysis of a single protein (i.e., the RdRp), such that we lack an understanding of genome-wide relationships and hence of the genesis and evolution of viral genera and species more broadly.

Glycoproteins are likely to be important determinants of major phenotypic characteristics across the *Flaviviridae*. They are essential for virus entry, influence host range and spillover potential, and are primary targets for host immune responses. However, glycoproteins have yet to be identified and/or classified for many species in the *Flaviviridae*. This cannot be resolved by even the most sensitive of sequence-based approaches ^10,13^, due to high levels of sequence divergence, and classical structural biology lacks the speed and scalability to sample enough species. This knowledge-gap limits investigation of molecular mechanisms, which in turn hinders the development of interventions such as vaccines.

We have augmented conventional phylogenetics with machine-learning-enabled protein structure prediction to comprehensively map glycoprotein structures across the *Flaviviridae*. This provided an evolutionary and genomic-scale perspective of the entire family, in doing so revealing molecular signatures that define the diverse virology and ecology found within the *Flaviviridae*.

## Results

### The Flaviviridae contains three major clades

Understanding the emergence and acquisition of molecular features across the *Flaviviridae* requires a proper gauge of the phylogenetic and genomic diversity of this family. To achieve this, we first constructed a comprehensive data set of flavivirus sequences: after clustering and manual curation this comprised 458 flavivirus genomes with complete coding sequences, including 11 that were novel taxa identified in this study (STable. 1). We next inferred a robust family-level phylogenetic tree for these data. Using conserved NS5 gene sequences that encode the RdRp, we applied various sequence alignment methods, quality trimming protocols, and amino acid substitution models, to infer a total of 225 phylogenetic trees for this family (STable. 2). Using distance-based approaches and manual inspection of the alignments and trees, we identified a phylogeny, denoted Tree 18 (STable. 2, SFig. 1, SFig. 2), that appeared to best represent the consensus topology of the *Flaviviridae*. Specifically, the topological placement of the major *Flaviviridae* clades in Tree 18 was consistent with 96% of phylogenies derived from MUSCLE and MAFFT multiple sequence alignments (MSAs), while this percentage dropped to 71% when Clustal Omega was included (see Methods for further details).

Our best-fit phylogenetic tree of the RdRp supported the division of the *Flaviviridae* into three distinct clades: (i) a clade comprising the large genome flaviviruses and members of the genus *Pestivirus*, (ii) an *Orthoflavivirus-Jingmenvirus* group, and (iii) a *Pegivirus-Hepacivirus* clade (Fig. 1A). Regardless of whether the tree was unrooted or rooted on a *Tombusviridae* outgroup, the LGF-*Pestivirus* and *Orthoflavivirus-Jingmenvirus* groups clustered together and formed a sister group to the *Pegivirus-Hepaciviru*s clade. The *Orthoflavivirus-Jingmenvirus* clade had the largest number of taxa (n = 182), followed by *Pegivirus-Hepacivirus* (n = 157) and LGF-*Pestivirus* (n = 119). All novel taxa fell within the LGF-*Pestivirus* clade.

**Figure 1.**
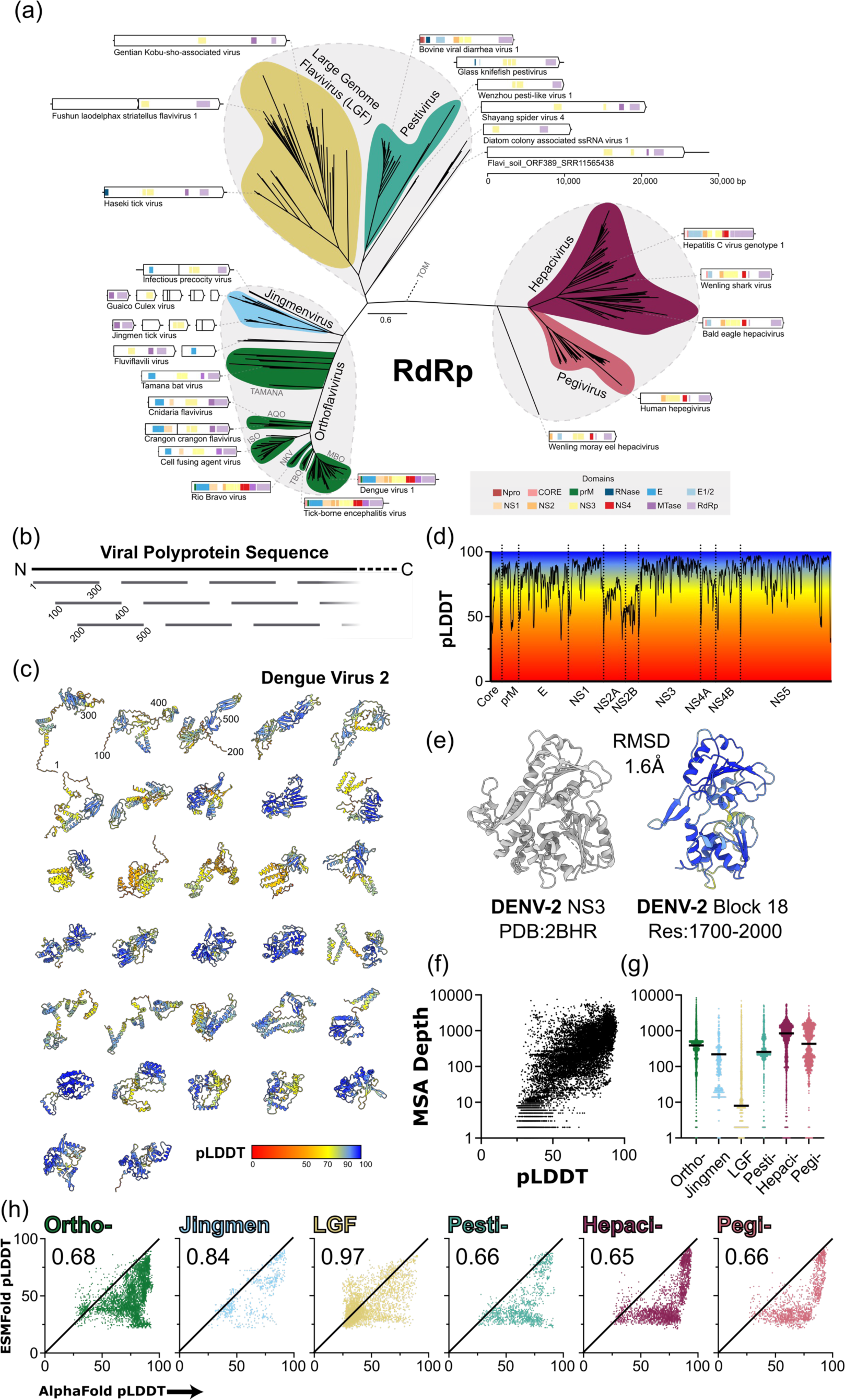
Generation of a Protein Foldome for the *Flaviviridae.* **A.** The RdRp phylogeny reveals three major lineages within the *Flaviviridae*: (i) *Orthoflavivirus*/*Jingmenmenvirus*, (ii) Large genome flavivirus/*Pestivirus* and (iii) *Hepacivirus*/*Pegivirus*. An unrooted tree is shown, with the *Tombusviruses* (TOM) representing the outgroup taxa and a scale bar denoting the number of amino acid substitutions per site. Genome organisation is provided for exemplar species, with annotations based on conserved domain sequence searches **B.** For protein structure prediction, all *Flaviviridae* polyprotein sequences from A were split into blocks of 300 residues, each overlapping by 100 residues (458 species, >16,000 blocks in total). Residue numbers are provided for the first three blocks. **C.** Representative AlphaFold protein structure predictions spanning the entire Dengue Virus 2 (DENV-2) polyprotein. Residue numbers are provided as in B. Structures are colour-coded by AlphaFold confidence scores (predicted local distance difference test; pLDDT), as denoted in the key. **D.** pLDDT confidence scores along the length of the DENV-2 polyprotein. Proteolytic cleavage sites delineating mature proteins are shown as dotted lines and protein names are provided on the x-axis. **E.** DENV-2 NS3 crystal structure (left) shown alongside an AlphaFold predicted structure for the corresponding region of the polyprotein (right). These structures superpose with a root mean square deviation (RMSD) of 1.6Å. **F.** Scatter plot demonstrating the relationship between multiple sequence alignment (MSA) depth and AlphaFold confidence (pLDDT). **G.** Distribution of MSA depths for each sequence block in each genus/subclade, colour-coded as in A. The mean is shown as a solid black line. **H.** Scatter plots representing the relationship between AlphaFold and ESMFold confidence for each sequence block in each genus/subclade. Numerical values are the performance ratio between either protein structure prediction method, with lower values indicating better performance in AlphaFold.

### ESMFold enables structural investigation of divergent species

We next aimed to explore protein functionality across the *Flaviviridae* using machine learning-enabled protein structure prediction. All flaviviruses encode polyproteins that undergo proteolytic maturation to liberate the constituent viral proteins. However, incomplete and ambiguous genome annotations, combined with extensive sequence divergence, make it very difficult to reliably identify the regions encoding each mature protein in all species. We therefore took a genome-agnostic approach, in which polyprotein coding sequences were split into sequentially overlapping 300 residue blocks for structural inference by two leading prediction models - AlphaFold and ESMFold (Fig. 1B & C) ^14,15^. This provided a comprehensive survey of protein structure across the *Flaviviridae* (458 species, >16K sequence blocks, >33K predicted structures), referred to here as the ‘protein foldome’.

As protein structural prediction has yet to be systematically applied in virology, we first evaluated folding performance. AlphaFold remains the most accurate prediction tool currently available and performed extremely well for many virus species. For example, ∼80% of the Dengue Virus 2 polyprotein yielded high confidence predictions (pLDDT > 70; Fig. 1C & D) and agreement with existing experimental structures (Fig. 1E). However, the accuracy of AlphaFold is directly proportional to the depth of the MSAs that guide structural inference. This was reflected in our data set, with shallow MSAs producing low confidence predictions (Fig. 1F). This becomes particularly problematic for the LGF, which are poorly sampled and, consequently, underrepresented in sequence databases, resulting in consistently shallow MSAs (Fig. 1G).

Structural inference by ESMFold is driven by a protein language model (pLM) and does not require MSAs, but is less accurate than AlphaFold. A comparison of folding confidence across our entire data set demonstrated that AlphaFold out-performed ESMFold in all viral genera with the exception of the LGF, in which they reached parity (Fig. 1H). This parity is not simply driven by poor performance from both methods; ESMFold yields informative predictions from sequences for which AlphaFold fails. Therefore, ESMFold, and other pLM-based approaches, may be particularly well suited for exploring the ‘viral dark matter’ revealed by metatranscriptomics.

### Discovery of glycoproteins across the Flaviviridae

Orthoflaviviruses, including Yellow Fever virus (the canonical species for the group), Tick-borne encephalitis virus (TBEV) and DENV, are often vector-borne. They possess the E glycoprotein; this represents a prototypical class-II fusion protein that undergoes a pH-dependent conformational cascade to merge viral and host membranes ^16^. Since the first discovery of class-II fusion proteins in TBEV ^17^, structurally and functionally homologous proteins have been identified outside of the *Flaviviridae*, both in viruses (e.g., Gc in the bunyaviruses ^18^) and in eukaryotes (e.g., HAP2 in plants, protists and invertebrates ^19^). The prevailing theory is that all instances of class-II fusion proteins share a common ancestor, although the origin of this progenitor remains unknown ^20^. In viruses, class-II fusion proteins are accompanied by a partner glycoprotein that is responsible for regulation and/or chaperoning of the fusogenic component. For example, in the orthoflaviviruses, E is partnered by the small prM glycoprotein ^21^. Partner proteins, however, are structurally diverse and may have multiple origins ^20^; for instance, prM is likely derived from host chaperonin proteins ^22^.

Identifying membrane fusion mechanisms in the hepaci-, pegi- and pestiviruses has proven more challenging. These viruses possess E1 and E2 glycoproteins that work in concert to achieve pH-dependent membrane fusion. The sequences of E1E2 bear no similarity to prM/E, and X-ray crystal structures of E2 from prototypical hepaci- and pestiviruses reveals folds that are broadly dissimilar from one another and from the E glycoprotein in orthoflaviviruses ^23–27^. Recent cryoEM analyses suggest that E1 adopts a unique fold, unlike that of any other known protein ^28,29^. Moreover, AlphaFold modelling, by ourselves, demonstrate that E1, unlike E2, adopts a highly conserved fold across the hepaci-, pegi- and pestiviruses, suggestive of mechanistic importance ^30^.

Whether E1E2 represents a novel and, as yet, uncharacterised fusion mechanism or a highly divergent iteration of a class-II system remains to be determined. Understanding the distribution, and biology, of glycoproteins across the *Flaviviridae* would likely provide insights on this. However, it is challenging to find or classify glycoprotein coding regions using sequence-based approaches in highly divergent species such as the jingmenviruses, LGF or basal members of the major subclades (for example, see incomplete sequence-based annotations in Fig. 1A). We overcame this limitation by performing structure similarity searches against the *Flaviviridae* protein foldome (Fig. 1). To achieve this, we built a custom reference library comprising experimental structures from the protein data bank (PDB) and published/validated AlphaFold models, and performed pairwise comparison with the entire foldome using a local installation of FoldSeek ^31^. To quantify structural similarity we used e-value outputs from the structure-guided sequence alignments performed by FoldSeek (low values indicating a high-probability match) and validated this approach using structures of SARS-CoV-2 spike and DENV NS5 RdRp as respective negative and positive controls (Fig. 2A & B).

**Figure 2.**
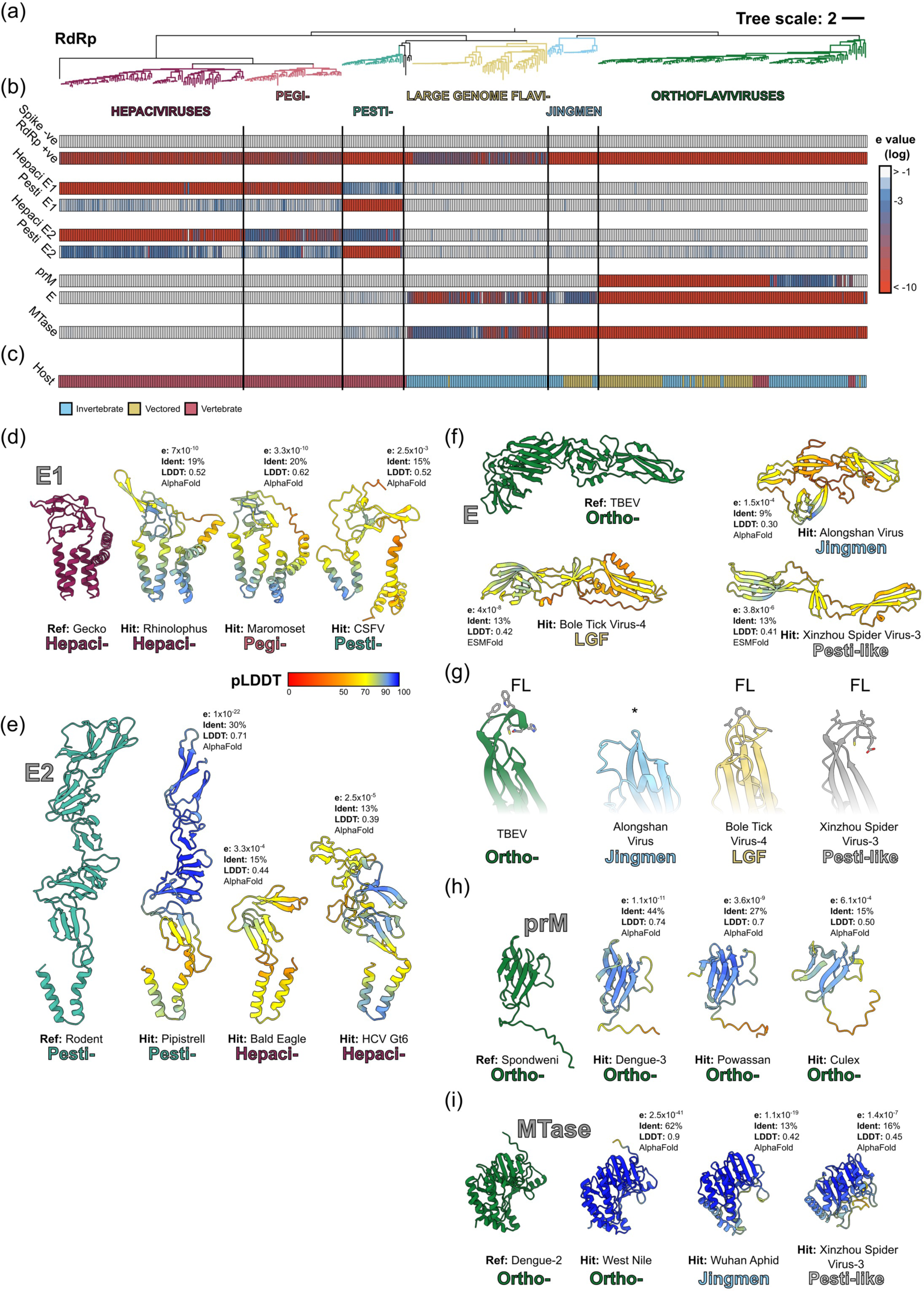
Discovery of glycoproteins across the *Flaviviridae*. **A.** RdRp phylogeny rooted on the tombusviruses, with each genus/subclade colour-coded as in Fig. 1A. **B.** FoldSeek e-value heatmaps for the indicated reference proteins, values are log transformed and colour-coded as shown in the key. For E1, E2, prM and E, the values represent summary e-values after comparison with a range of relevant reference structures, as described in the methods. **C.** Host species tropism for each virus, where “Vectored” refers to those assigned as “Yes” or “Potentially” in STable. 3. **D-I.** Representative reference structures and FoldSeek hits for E1, E2, E, prM and methyl transferase (MTase). For each hit only the FoldSeek aligned residues are shown for any given sequence block, metrics provide e-value, sequence identity (%), structural alignment score (local distance difference test, LDDT, ranging from 0 to 1), and protein structure prediction method. Predicted structures are colour-coded by pLDDT confidence scores, as shown in the key. For D. and E., the reference structures are previously published AlphaFold models, for F., G. and I. experimental structures are used (PDB:6ZQI, 7QRF, 1L9K respectively). **H.** Structurally conserved E protein fusion loops (FL) found in orthoflavi-, LGF-, and in pesti-like viruses. The FL is absent from the E protein homologue of the jingmenviruses (its expected location marked by an asterisk). Amino acid side chains are shown for the FL only. CSFV, Classical swine fever virus; HCV, hepatitis C virus; TBEV, Tick-borne encephalitis virus.

The FoldSeek e-value metric was sufficiently sensitive to detect the high structural homology between hepaci-/pegi- and pestivirus E1, despite sharing only 10-15% sequence identity (Fig. 2B & D). Hepaci-/pegi- and pestivirus E2 are structurally divergent ^30^, nonetheless FoldSeek demonstrated clear reciprocal structural similarity focussed on the C-terminal portion of E2 (proximal to the transmembrane domain) where sequence identity ranged from 8.5-15% (Fig. 2B & E). The distribution of E1 and E2 were in near-perfect correlation, consistent with mechanistic interdependence, and we found no evidence for E1E2-like folds outside of the hepaci-/pegi- and pestiviruses.

Using DENV and TBEV as references, we mapped structural homologues of E glycoprotein to the orthoflaviviruses, jingmenviruses, LGF and pesti-like species that sit basal to the classical *Pestivirus* subclade (e.g., Xinzhou spider virus 3) (Fig. 2B & F). Notably, successful mapping was dependent on ESMFold predicted structures for the most divergent sequences (LGF and pesti-like), emphasising the value of using complementary prediction methods (Fig. 2F). A notable exception was a group of viruses of unknown hosts discovered in environmental samples ^32–34^ (e.g., Inner Mongolia sediment flavi-like virus 3) for which no glycoproteins were identified (SFig. 3); whether these represent species without structural proteins or partial genomes remains to be determined.

For most E glycoprotein homologues, the structural prediction was sufficient to identify the hydrophobic fusion loop at the tip of domain II, which inserts into host membranes and is central to the class-II fusion mechanism (Fig. 2G). However, the fusion loop was absent from the jingmenvirus E homologues, suggesting significant mechanistic divergence in these viruses, likely coincident with their unique genome segmentation. In contrast to E glycoprotein, we were only able to detect the prM partner glycoprotein within the orthoflaviviruses (Fig. 2B & H). A critical function of prM is occluding the fusion loop of E during particle maturation and we may yet expect to find orthologous partners in other clades.

### Glycoprotein distribution correlates with ecological niche

We could unambiguously assign either E1E2 or E glycoproteins to the vast majority of species in the *Flaviviridae* (Fig. 2B). Their distributions broadly divide the family in two, although this division is incongruent with RdRp phylogeny suggesting a complex evolutionary history; in particular, the pestiviruses and LGF, that represent sister clades, possess E1E2 and E, respectively. While mapping glycoproteins was the primary focus of our study, we also compared the foldome against any *Flaviviridae* proteins with entries on the PDB. In doing so we observed that all species with a methyltransferase (MTase) also possess an E glycoprotein homologue (Fig. 2B & I). Viruses with MTase undergo cap-dependent translation, whereas those without (hepaci-, pegi-, pestiviruses) rely on an internal ribosome entry site (IRES). We reasoned that E/MTase and E1E2/IRES may represent divergent co-adaptations to particular ecological niches and, to assess this, we compared virus-host associations across the phylogeny (Fig. 2C). Viruses with E/MTase infect a variety of hosts, including those that are transmitted between vertebrates by invertebrate vectors (e.g., DENV by *Aedes* mosquitoes). In contrast, E1E2/IRES (i.e., lack of MTase) was strictly correlated with vertebrate hosts. This suggests that the gain of E1E2 and an IRES, with the concomitant loss of MTase, represent a molecular commitment to replication and transmission in vertebrates. Moreover, based on the underlying RdRp phylogeny, this commitment to vertebrates is likely to have occurred twice in the *Flaviviridae*, once for the hepaci-/pegiviruses and once for the pestiviruses.

### The E and E1E2 glycoproteins are evolutionarily distinct

Through systematic protein structure prediction we have revealed deep homology that cannot be detected using sequence-based approaches. To further explore these relationships we performed phylogenetic analyses of structurally aligned sequences, derived from FoldSeek. For the E glycoprotein, this results in a phylogeny that is largely reflective of the RdRp tree, with E homologues from orthoflavi-, jingmen- and LGF forming three distinct clades (Fig. 3A). This is consistent with evolution from a common ancestral virus that possessed a class-II fusion mechanism. Notably, the E protein homologues in pesti-like viruses of spiders cluster within the LGF glycoprotein clade (Fig. 3A), similar to the RdRp tree topology (Fig. 1A & 2A). This suggests that the gain of E1E2, accompanied by loss of E, was a defining event in the emergence of the pestiviruses from an LGF-like progenitor (Fig. 3B).

**Figure 3.**
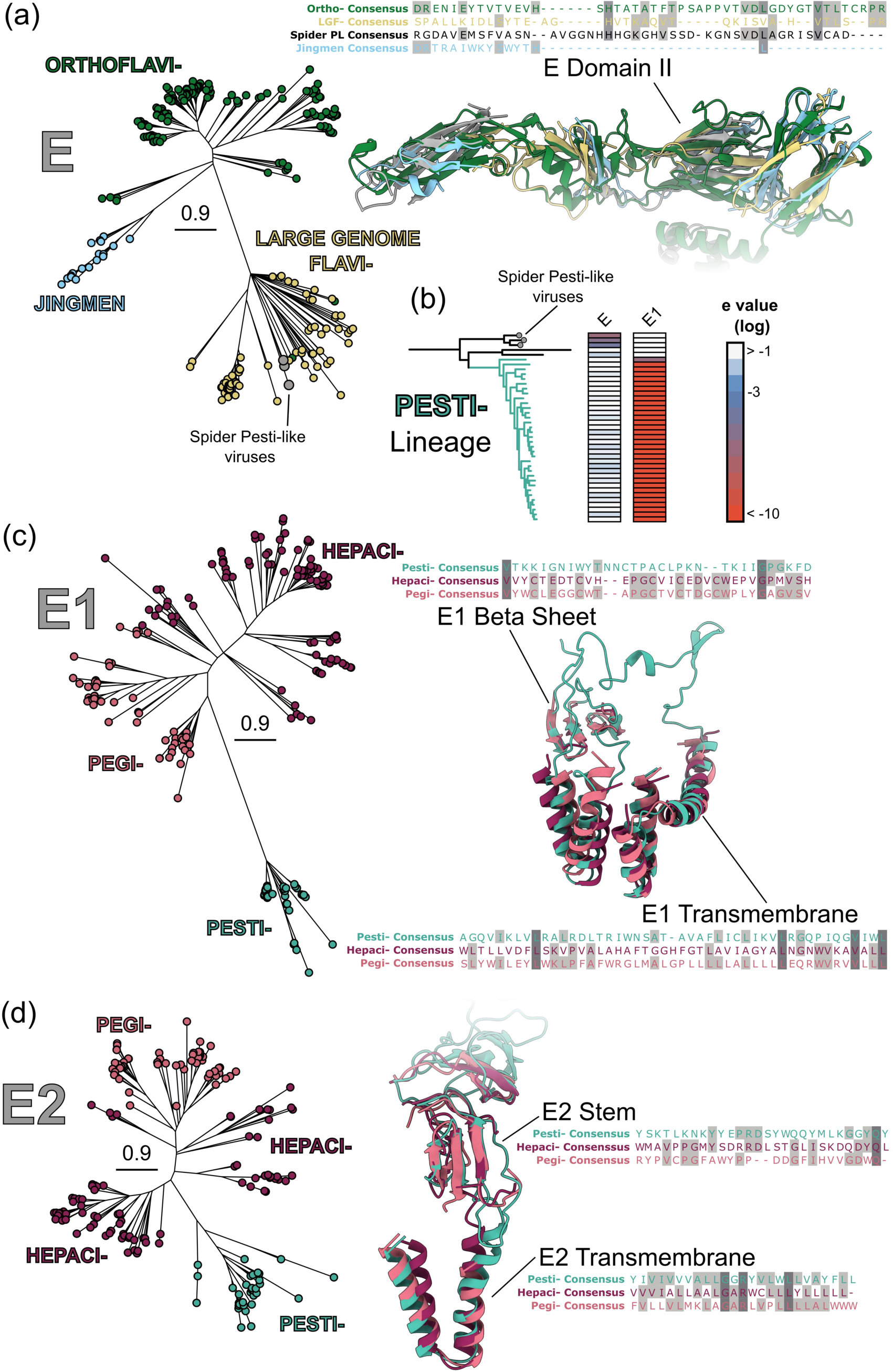
Structure-guided phylogenies of E, E1 and E2 glycoproteins. **A.** E protein phylogenies were constructed using structure-guided sequence alignments from FoldSeek, with the scale bar indicating the number of amino acid substitutions per site. The right displays representative superposed structural elements. Structures were aligned using flexible FATCAT with a DENV1 E protein AlphaFold model as reference (shown in green). The Spider pesti-like virus E glycoprotein is shown in grey, other structures are colour-coded as in the phylogeny. The associated protein alignment provides an example of a structurally aligned region (domain II of E), and contains consensus E glycoprotein sequences for the stated genus/subclade with conserved sites highlighted in grey. **B.** *Pestivirus* sublineage RdRp phylogeny and FoldSeek e-value heatmaps for the stated reference structures. The spider pesti-like viruses basal to the pestiviruses possess E protein and not E1 (or E2). **C.** Structural phylogeny and example structural alignments for E1. Flexible FATCAT alignments were performed using the BVDV E1 AlphaFold model as reference. Structures are colour-coded as in phylogeny **D.** Structural phylogeny and example structural alignments as for E2, flexible FATCAT alignments were performed using Rodent pestivirus E2 AlphaFold model as reference. Here structural/sequence similarity was focussed on the basal region of E2, in particular the stem and transmembrane domains.

Both E1 and E2 alignments and phylogenies suggest a common glycoprotein ancestry in the hepaci-, pegi- and pestiviruses (Fig. 3C & D), despite the pesti- and hepaci/pegi lineages appearing paraphyletic in the RdRp phylogeny (Fig. 1A). We could not detect any significant structural homology between E1E2 and E, or identify intermediate forms between these glycoprotein systems, further suggesting they are mechanistically distinct. Therefore, based on current evidence we propose that E1E2 represents a novel class of fusion protein that potentially emerged in the *Hepaci*-/*Pegivirus* lineage and underwent genetic transfer to an LGF-like virus to give rise to the pestiviruses.

### Novel and acquired proteins in a large genome flavivirus

Our structure-guided approach is particularly suitable for gaining insights into divergent and/or poorly characterised viruses such as those found in the LGF. Whilst the majority of LGF species likely infect invertebrates, there is evidence that one subclade, the bole tick viruses, are capable of tick-borne infection of mammals including humans ^3^. This group may represent an emergent threat to public health and, therefore, warrants closer scrutiny.

Focusing on Bole Tick Virus 4 (BTV4), we examined the N-terminal portion of the polyprotein proximal to the E glycoprotein homologue identified in our initial analyses (Fig. 2). The N-terminal structural proteins of *Flaviviridae* polyproteins are typically processed by host signal peptidases to liberate the mature proteins. Therefore, instead of arbitrary 300 residue sequence blocks, we used cleavage site prediction to identify five putative protein coding sequences (labelled A-E, Fig. 4A). Protein structures were predicted using three approaches - ESMFold, AlphaFold and AlphaFold with manually-curated MSAs - with the latter only possible due to our comprehensive survey and phylogenetic analysis (Fig. 1A). For each sequence, we provide the highest confidence model produced by any of these methods (Fig. 3A).

**Figure 4.**
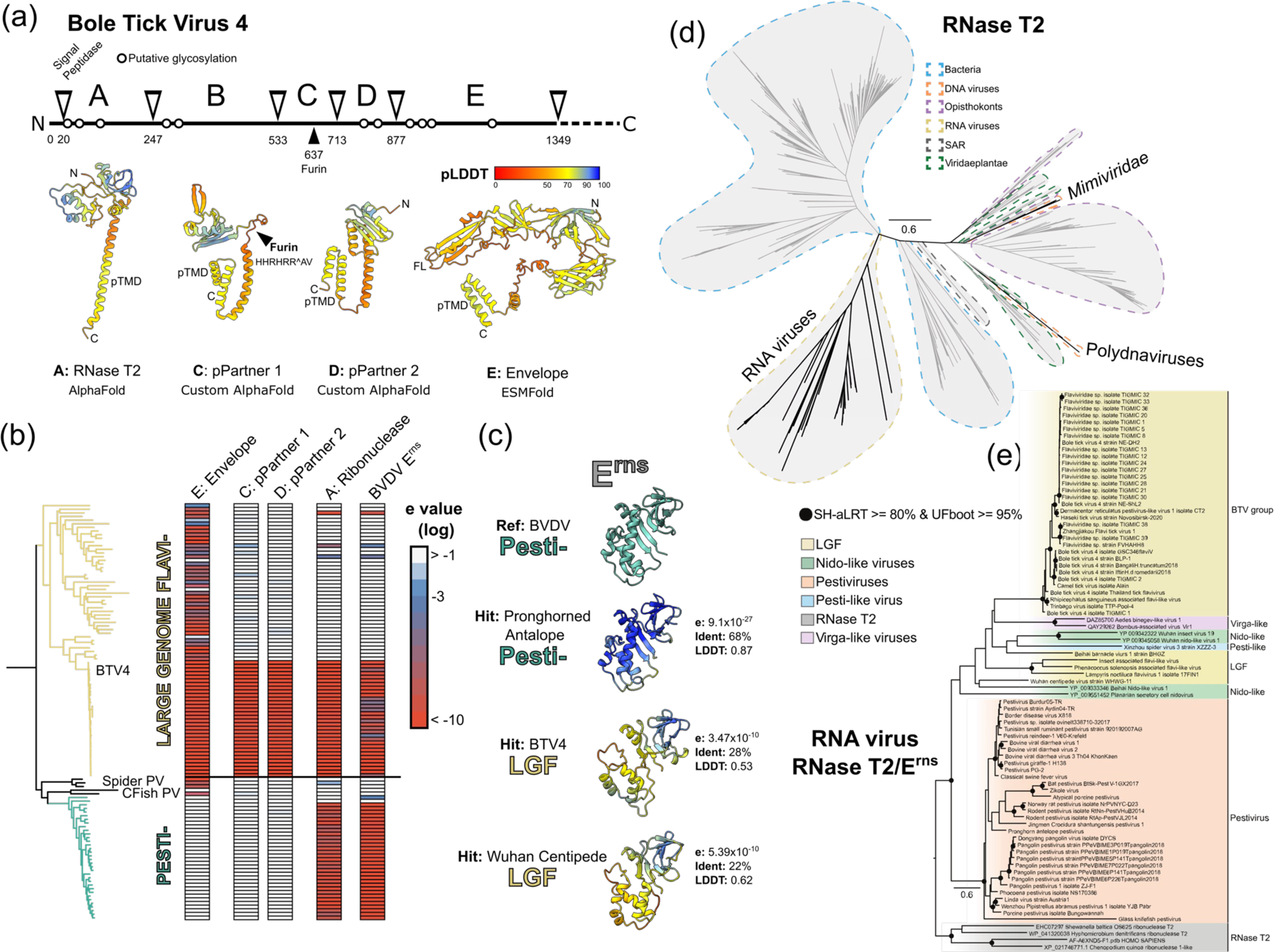
Novel and acquired proteins in a large genome flavivirus. **A.** N-terminal glycoproteins from Bole Tick Virus 4 (BTV4). Linear representation of the BTV4 polyprotein displays location of putative glycosylation and the predicted signal peptidase cleavage sites that delineate five mature proteins (labelled A-E). For each of these mature proteins, highest confidence models are shown from three prediction methods (AlphaFold, AlphaFold with custom MSAs and ESMFold). Protein B did not yield any high-confidence model and is not shown. The inferred functions/names of the proteins are as described in the main text. Each protein contains a putative transmembrane domain (pTMD), protein C contains a canonical furin cleavage site, the conserved fusion loop (FL) of E is also annotated. **B.** Large genome flavivirus/*Pestivirus* lineage RdRp phylogeny and FoldSeek e-value heatmaps for the stated reference structures, annotations provide the location of BTV4 (reference), the spider pesti-like viruses (Spider PV), and the cartilaginous fish pesti-like viruses (CFish PV). **C.** Example FoldSeek hits against an experimental structure of bovine viral diarrhoea virus E^rns^ ribonuclease (PDB:4DVK). **D.** Ribonuclease T2 sequence phylogeny, with domains of life/viruses colour-coded as shown in the key, scale bar indicates phylogenetic distance (substitutions/site). This protein has been independently acquired once by RNA viruses and twice by DNA viruses (in *Mimiviridae* and polydnaviruses). The RNA virus clade is nested within bacterial instances of the ribonuclease T2, suggesting a single horizontal gene transfer event. **E.** Phylogeny of the RNA virus RNase T2/E^rns^ clade rooted on non-viral sequences, with viral clades colour-coded as shown in the key. BVDV, Bovine viral diarrhoea virus.

The largest and most C-terminal protein is the E glycoprotein homologue. ESMFold produces a model of intermediate confidence in which the transmembrane domain and fusion loop are juxtaposed; this is consistent with experimental structures of the post-fusion conformation of E ^35^. Directly upstream of the E homologue are two smaller proteins for which custom AlphaFold yielded the optimal predictions. Whilst neither protein shares direct homology with prM, they both have a broadly similar organisation, consisting of a small globular domain anchored by a putative transmembrane region. Therefore, given their direct proximity to the E homologue, we suggest that these are partner proteins that provide chaperoning to the class-II fusogen. Indeed the putative partner 1 possesses a furin cleavage site in-between its globular domain and membrane anchor. This is completely analogous to prM, and would provide a means of proteolysis during secretion, akin to the maturation of orthoflavivirus particles ^21^. Protein coding sequence B yielded low confidence predictions from each folding approach and is not shown. The most N-terminal sequence produced a good AlphaFold model, which structural similarity searches identified as a T2 family ribonuclease (RNase T2) with homologs across the tree of life.

We used FoldSeek to investigate the distribution of these protein structures across the *Pestivirus*/LGF clade (Fig. 3B). Consistent with orthoflavivirus E glycoprotein references (Fig. 2), homologues of BTV4 E glycoprotein were detected throughout the LGF and in pesti-like viruses identified in spiders and cartilaginous fish that fall basal to members of the classical genus *Pestivirus*. Therefore, using a proximal reference (BTV4 E glycoprotein), we provide further evidence for the loss of E, and gain of E1E2, at the genesis of the pestiviruses. In contrast, the BTV4 putative partner proteins were confined to the Bole Tick Virus subclade, and structural similarity searches against current protein databases (e.g. PDB and AlphaFoldDB) revealed no homologues. Therefore, these proteins are likely novel and may represent an adaptive feature, specific to these viruses.

BTV4 RNase T2 has homologues throughout the Bole Tick Virus subclade, in some other LGF species and, notably, across the genus *Pestivirus*, where the homology maps to the E^rns^ ribonuclease. Indeed, FoldSeek searches using an E^rns^ crystal structure ^36^ reveal a reciprocal distribution of homology (Fig. 3B & C). Phylogenetically, the pesti-/LGF E^rns^ form a deep branch amongst homologous RNase T2 sequences from viruses, bacteria, plants and animals (Fig. 3D & E). Together, this indicates that E^rns^ originated in a distant ancestor of the pesti- and LGFs, likely from a single horizontal gene transfer of a bacterial RNase T2 giving rise to all extant instances of *Flaviviridae* E^rns^. Moreover, the distribution of E^rns^ is broadly concordant with RdRp phylogeny, suggesting that E^rns^ has been continuously retained in certain species and lost in others (Fig. 3E), as opposed to undergoing genetic exchange through recombination within the clade. Further instances of RNase T2 from nido- and virga-like viruses were also nested within the E^rns^ tree, indicating onward genetic transfer to other RNA viruses.

## Discussion

By mapping the distribution of glycoprotein structures we gained insights into the deep evolutionary history of the *Flaviviridae*. Specifically, we propose the existence of two distinct evolutionary lineages, as summarised in Fig. 5. Lineage one, encompassing the *Orthoflavivirus/Jingmenvirus* and LGF/*Pestivirus* clades, arose from an ancestor that possessed the E glycoprotein and performed cap-dependent translation, necessitating MTase. Lineage two contains the *Hepacivirus/Pegivirus* clade and arose from an ancestor that possessed E1E2 glycoproteins and lacked MTase, implying reliance on IRES-dependent translation. The characteristics of the common ancestor of these two lineages (i.e., the common ancestor of the *Flaviviridae*) remain speculative. Based on the taxonomic distribution of infected hosts and the existence of endogenous viral elements, it has been suggested that this common ancestor may have originated over 900 million years ago ^37^. Our evolutionary reconstruction suggests that this ancestor possibly contained core NS3/NS5 proteins and potentially an MTase. Although the absence of RrmJ-like methyltransferases, such as those found in the *Flaviviridae,* in other Kitrinoviricota members^38^, hints that MTase might have been acquired at the base of lineage one. It remains unclear whether this progenitor possessed a viral envelope, therefore necessitating entry glycoproteins, although it is noteworthy that all Kitrinoviricota members (n = 20) with the exception of *Togaviridae* and *Matonaviridae* are non-enveloped.

**Figure 5.**
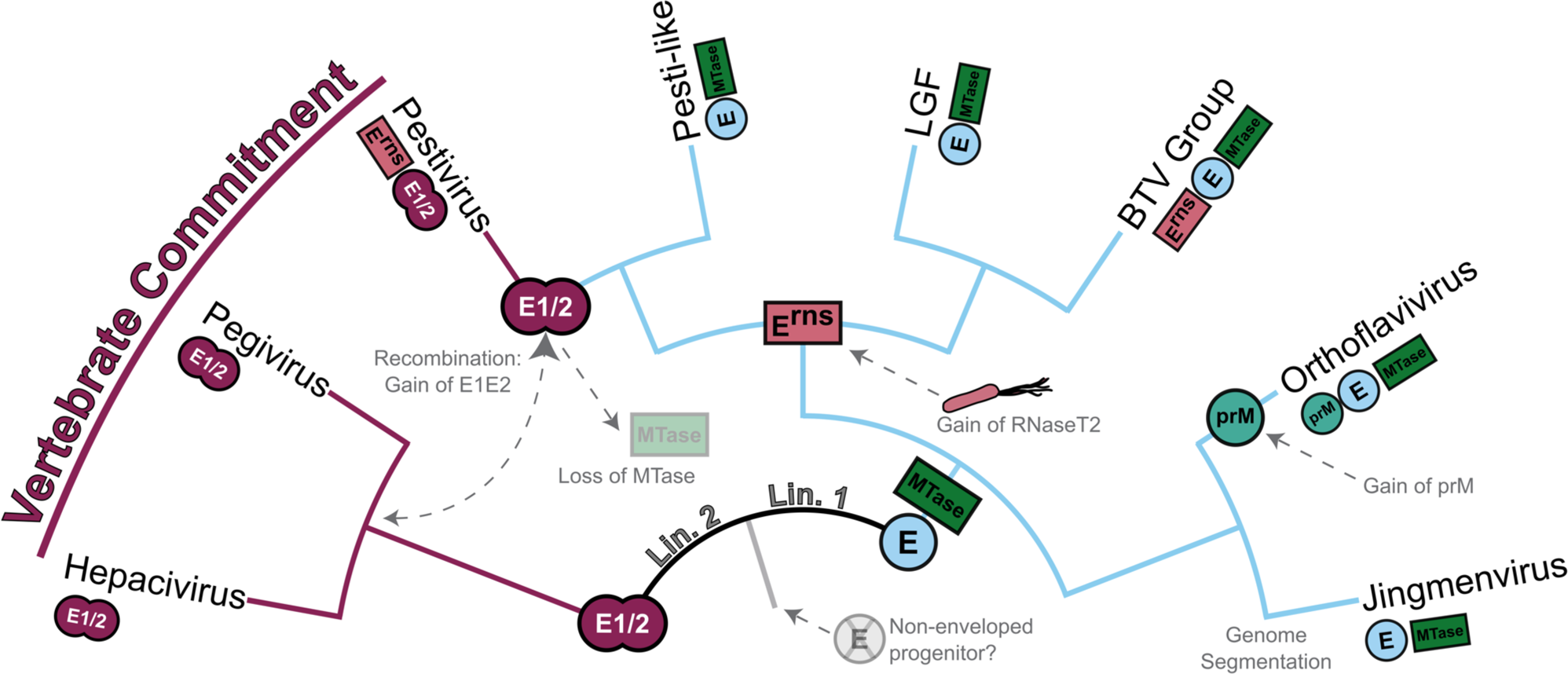
Proposed evolutionary history of the *Flaviviridae*. Illustrative cladogram showing the key protein acquisition and loss events across the major *Flaviviridae* clades. The two major lineages, described in the main text, are labelled (Lin. 1, Lin. 2), near the root. Each clade, represented as a tip, is annotated with symbols representing the presence of key proteins. Branches are highlighted to denote the lineage-specific presence of envelope protein E (in light blue) or E1/E2 (in maroon). Major nodes are emphasised with larger symbols to infer the ancestral emergence of each protein within the evolutionary history of the *Flaviviridae*. Dashed lines and arrows denote the loss or gain of specific proteins, highlighting potential recombination events and gene transfers that have shaped the *Flaviviridae*.

Lineage one has undergone extensive diversification. Genome segmentation occurred in the jingmenviruses, with coincident divergence in their E glycoprotein ^13^, including the apparent loss of its fusion loop. The orthoflaviviruses gained prM, a partner to the E glycoprotein, derived from a host chaperonin ^22^. The sister lineage gives rise to the LGF and *Pestivirus* subclades. Here, an ancestral virus gained RNase T2, likely from a bacterial lineage, which was retained both in the Bole Tick Virus group of the LGF and in the pestiviruses. The phylogeny of pestivirus/LGF RNase T2 is broadly consistent with that of the cognate RdRp and hence further supports a shared evolutionary history and a single origin. Moreover, in both cases RNase T2 is found at the N-terminus of the polyprotein juxtaposed with the glycoproteins (e.g., E^rns^ directly precedes E1 in the pestivirus genome). We were able to unambiguously identify the E glycoprotein in the majority of LGFs, and in pesti-like viruses that sit basal to the classical pestiviruses. However, as all classical pestiviruses possess E1E2 glycoproteins, homologous to those found in the hepaci-/pegiviruses; there was seemingly a switch in glycoprotein systems through an inter-lineage recombination event.

In contrast, lineage two exhibits relatively little diversity, with the hepaciviruses and pegiviruses sharing similar genome organisation, and relatively high conservation in the sequence and structure of RdRp and E1E2. However, we anticipate that considerable diversity exists along the ancestral *Hepacivirus*/*Pegivirus* branch reflecting both extinct viruses and extant species that remain to be discovered.

The presence of E1E2 is strictly associated with viruses of vertebrates. This suggests that E1E2-dependent virus entry represents a molecular commitment to the vertebrate niche. This commitment also appears to extend to species and, even, tissue tropism. For instance, HCV can only efficiently enter human hepatocytes, with E1E2 activity requiring a very specific set of entry factors ^39^. Indeed, hepatotropism is likely a feature of all hepaciviruses as they are frequently discovered and abundant in the liver tissue of mammals, lizards and fish ^40–42^. Therefore, glycoprotein biology may have locked these viruses into narrow ecological niches, which may explain the comparative lack of divergence in the *Hepacivirus*/*Pegivirus* lineage.

Conversely, the class II fusion protein fold, exemplified by the E glycoprotein, is found in viruses with a wide variety of hosts, particularly in the *Flaviviridae* where it spans the breadth of the Metazoa. This protein is also found in other viral families that are vectored from invertebrates to vertebrates and, in some cases, plants ^20^. Additionally, Orthoflaviviruses exhibit a wide range of tissue tropisms within a single host ^43^. This suggests that the class II mechanism possesses relatively high versatility in its biochemical/biophysical requirements for achieving membrane fusion. This may permit lower host stringency, facilitating adaptation to diverse niches and transmission cycles. This is consistent with both the divergence of *Flaviviridae* lineage one and the ongoing zoonotic threat posed by viruses in the orthoflaviviruses, jingmenviruses and LGFs.

We observed no structural similarity between the E and E1E2 glycoproteins. This is unlikely to be attributable to a lack of sensitivity; we detected strong structural homology between E glycoprotein homologues that were >85% divergent at the sequence level. This suggests that E1E2 represents a novel membrane fusion mechanism that arose de novo in the *Flaviviridae*. This may have occurred on the long, and unsampled, branch leading to the hepaci-/pegiviruses, before undergoing recombination with an LGF-like virus to give rise to the pestiviruses. Considering the phylogenetic placement of the arachnid-associated pesti-like viruses, it is tempting to speculate that the pestiviruses originated from arachnid viruses and subsequently acquired E1/E2 through recombination, possibly within a blood-feeding host. Indeed, a similar scenario involving hematophagy in ticks has been suggested for the origins of vertebrate infection in the orthoflaviviruses ^37^. However, as yet, there is insufficient evidence to confidently infer the directionality of E1/E2 genetic transfer, or its origins.

In sum, we demonstrate that machine-learning-enabled protein structure prediction can provide novel insights and clarity into the evolutionary history of viruses. Whilst AlphaFold offers unparalleled prediction confidence and accuracy, pLM-based systems, such as ESMFold, may be more capable in the analysis of highly divergent species at the fringes of the known virosphere. Indeed, ESMFold analysis of the LGF not only identified the viral glycoproteins, but also an apparent acquisition of a bacterial enzyme (RNase T2). Using the same approach, we recently discovered other apparent horizontal transfers from prokaryotes to an LGF-like virus with a 39.8kb genome ^10^. It is therefore possible that these viruses exhibit frequent “genetic piracy” and that the high coding capacity, afforded by their long genomes, plays host to a diverse range of acquired functions. Through generating the *Flaviviridae* foldome data set alongside the curated complete genome and phylogeny set in the current study, we offer a valuable resource for future flavivirus research. Our study highlights how protein structure prediction represents the new state-of-the-art tool for disentangling the complex evolutionary-scale relationships between viruses and the broader tree of life.

## Acknowledgments

We acknowledge the University of Sydney’s high-performance computing cluster, Artemis, for providing the computing resources used for this study. E.C.H. is supported by a National Health and Medical Research Council Investigator award (GNT2017197). J.C.O.M. is supported by the Australian Government’s Research Training Program Scholarship. J.G., S.L., and M.O. are supported by the Wellcome Trust and Royal Society through a Sir Henry Dale Fellowship (107653/Z/15/Z). K.T. was supported by a Lord Kelvin Adam Smith Fellowship from the University of Glasgow. J.G. is also supported by the MRC-University of Glasgow Centre for Virus Research core support from the Medical Research Council/UKRI (MC_UU_00034/1).

## Author contributions

Conceptualization, J.C.O.M., E.C.H., and J.G.; Methodology, J.C.O.M., S.L., M.R.O., K.T., E.C.H., and J.G.; Formal Analysis, J.C.O.M., S.L. and J.G.; Investigation, J.C.O.M., S.L., V.A.C., and J.G; Visualization, J.C.O.M., S.L. and J.G.; Supervision, E.C.H. and J.G.; Funding Acquisition, E.C.H. and J.G.; Writing - Original Draft, J.C.O.M. and J.G. Writing— Review and Editing, all authors.

## Data availability

All underlying data, including sequences and structures, are available here: https://zenodo.org/doi/10.5281/zenodo.10616318

## Methods

### Compilation of Flaviviridae sequence set

#### Retrieval of flavivirus genomes

Flavivirus sequences were collected using the search phrase “*Flaviviridae* taxid 11050 and Unclassified *Flaviviridae* taxid 38144” in the NCBI Virus Database on the 15^th^ of December 2022. The search was complemented by referencing sequences from Mifsud et al. 2022 ^42^ and supplemented with sequences from the NCBI nucleotide database using the search phrase “flavi[All Fields] OR pesti[All Fields] OR hepaci[All Fields] OR pegi[All Fields] AND viruses[filter]” on the same date. Additional sequences were later retrieved from publications that had sequences not available in GenBank at the time ^9,33,44–49^.

#### Sequence set curation

Sequences were clustered to a 95% nucleotide identity threshold to approximate a species-level distinction, excluding the LGF tick-associated clade. Clustering was performed using CD-HIT (v4.6.1) ^50^ with non-default parameters “cd-hit-est -c 0.95 -n 9”. Subsequently, the clustered sequence set was manually curated by removing incomplete coding regions. Sequences shorter than 2,000 nucleotides in length were removed, with the exception of the jingmenviruses where segments are known to be <2000 nucleotides in length. These nucleotide sequences were translated using the Geneious Prime Find ORFs tool (v2022.0) (www.geneious.com) ^51^ and along with protein sequences aligned to annotated reference sequences (where available) using MAFFT FFT-NS-I X2 (v7.402) to assess genome completeness ^52^. This was complemented by predicting conserved domains using the InterProScan software package (v5.63-95.0) with the SFLD (v4.0), PANTHER (v17.0), SuperFamily (v1.75), PROSITE (v2022_05), CDD (v3.20), Pfam (v35.0), SMART (v9.0), PRINTS (v42.0), and CATH-Gene3D databases (v4.3.0) (Jones et al. 2014). Sequences determined to contain partial coding sequences were removed from the subsequent analyses.

#### Discovery of novel LGF sequences

Tick-associated LGFs are of particular interest due to the recently reported association between Haseki tick virus and tick-borne infectious disease in humans (Kartashov et al., 2023). To identify related viruses, we screened the Sequence Read Archive (SRA) RdRp microassemblies generated by Serratus ^53^ using DIAMOND BLASTx (v2.0.9) ^54^ (e-value threshold of 1E^−5^ and the “--ultra-sensitive” flag) (Buchfink et al., 2021) with Haseki tick virus (UTQ11742) as the query. An e-value threshold of 1.6E^−1^ was established to restrict the number of libraries for reassembly to a manageable quantity. This threshold was determined based on the organism associated with the SRA library and the percent identity values. The 319 SRA libraries that meet this threshold were processed following the BatchArtemisSRAMiner pipeline ^55^. Briefly raw FASTQ files were retrieved using Kingfisher (https://github.com/wwood/kingfisher-download), quality trimming and adapter removal using Trimmomatic (v0.38) (Bolger et al. 2014) with parameters SLIDINGWINDOW:4:5, LEADING:5, TRAILING:5, and MINLEN:25 and de novo assembly using MEGAHIT (v1.2.9) ^56^ with default parameters. The assembled contigs were compared to the NCBI non-redundant protein database (as of March, 2023) and a custom *Flaviviridae* protein database using DIAMOND BLASTx as described above. All novel flaviviruses predicted to contain complete coding sequences identified by this method (including those outside of the LGF group) were included in phylogenetic analyses.

### Protein structure prediction and homology search

#### Systematic protein structure prediction

We adopted a strategy to overcome incomplete and ambiguous genome annotations, and generate sequence lengths that are amenable to rapid inference of structure. *Flaviviridae* polyprotein amino acid sequences were broken into sequential 300 residue blocks with a 100 residue overlap. However, most polyproteins are not equally divisible by 300, therefore, we set the final sequence block to cover the final 300 residues of the polyprotein, irrespective of overlap with the penultimate block. This resulted in 16,619 sequence blocks from 561 species (558 from the *Flaviviridae* and 3 from the *Tombusvirus* outgroup). Structures were predicted for each sequence using the ColabFold implementation of AlphaFold, with default settings but only generating a single model per target, performed using Google Colab cloud computing. Structural inference was also performed with ESMFold (using the 3 billion parameter ESM-2 model), on local compute (Nvidia V100 GPU + 32GB vRAM). This resulted in a total of 33,238 structural models and associated metadata (e.g ColabFold MSAs). Custom Python scripts were used to break up sequences for folding and extract metrics from outputs (i.e. pLDDT confidence and MSA depth). All structural data was prepared for publication using UCSF ChimeraX ^57^. Representative structural superpositions (Fig. 3) were performed using FatCat 2.0 ^58^. For inference of putative mature protein sequences (Fig. 4) the SignalP server was used to predict the junctions between viral proteins ^59^. For custom AlphaFold inference (Fig. 4), whole polyprotein sequences of the Bole Tick Virus group were aligned using MAFFT, MUSCLE (v5.1) ^60^, and subalignments covering only the putative protein sequences were converted to the .a3m format and used as input for ColabFold structure prediction ^61^. All predicted structures are included in the underlying data.

#### Structural homology searches

We used FoldSeek in exhaustive search mode to cross compare the *Flaviviridae* protein foldome with a library of reference structures drawn from the protein database and AlphaFold models of particular targets. FoldSeek was set to output e-values, structurally aligned amino acid sequences, % identity of aligned residues, bit score and lddt structural similarity, with an e-value cut off of 0.1 to limit the size of the output datafile. To interrogate the output data, the lowest e-value scores for any given species against any given reference structure were extracted using a custom python script. Where multiple references were used for a single protein (e.g., E glycoprotein, 3 references: PDB:6ZQI, PDB:7QRF, AlphaFold model: DENV1 E) the lowest e-value against any given species was chosen. This data was plotted against sequence-based phylogenies using the Interactive Tree Of Life ^62^.

Representative hits (Fig. 2D-I) were selected manually to reflect the levels of similarity and divergence in structure and sequence. All reference structures are included in the underlying data, the following experimental structures, from the PDB, were used: 6ZQI, 1L9K, 5F3Z, 7QRF, 7V1E, 7T6X, 6VYB, 2YQ2 and 4DVK ^22,23,28,36,63–67^.

### Phylogenetic analysis

#### NS5b phylogeny

The evolutionary relationships among the *Flaviviridae* were inferred using maximum likelihood (ML) phylogenies derived from MSAs of the highly conserved NS5b region (that encodes the RdRp). This region was extracted from each sequence by aligning polyprotein sequence subsets according to their taxonomy and using both pre-existing and newly generated NS5b annotations from InterProScan as a guide. As alignment and trimming parameters have been shown to influence the topology of the *Flaviviridae* ^68^ we compared several methods resulting in 225 phylogenies. Briefly, flavivirus sequences were aligned using MAFFT, MUSCLE (v5.1) ^60^ and Clustal Omega (v1.2.4) ^69^ with default parameters. Ambiguously aligned regions were removed using trimAl (v1.2) ^70^ with eight conservation thresholds (i.e., minimum percentage of alignment columns to retain) - 5, 7.5, 10, 12.5, 15, 17.5, 20, and 25 - and three gap thresholds (i.e., the minimum fraction of sequences without a gap needed to keep a column) - 0.7, 0.8, and 0.9 - as well as the automated parameter selection method “gappyout”.

All ML phylogenetic trees were estimated using IQ-TREE 2 (v1.6.12) ^71^. Selection of the best-fit model of amino acid substitution was inferred for a subset of phylogenies using the ModelFinder function in IQ-TREE 2 ^72^. In addition to the model chosen by ModelFinder (LG+F+R10) two additional models, the Le-Gascual model (LG) and FLAVI ^73^ were compared. Branch support was calculated using 1,000 bootstrap replicates with the UFBoot2 algorithm and an implementation of the SH-like approximate likelihood ratio test within IQ-TREE 2 ^74,75^. To root the phylogeny, three members of the *Tombusviridae* family were chosen given their remote sequence similarity to the NS5 region of the *Flaviviridae* ^7,76^. Phylogenetic trees were annotated using the R packages phytools (v1.5-1) ^77^, and ggtree (v3.3.0.9) ^78^ and further edited in Adobe Illustrator (https://www.adobe.com). Genome diagrams were constructed using a manually curated selection of predicted functional domains and visualised using gggenomes (v0.9.8.9) ^79^.

For each virus sequence, host information was pulled from the corresponding GenBank “host” field using rentrez (v1.2.3) ^80^ and standardised using taxize (v0.9.1) ^81^. Vector status, defined as “yes”, “no”, and “potentially”, was assigned by first querying the Arbovirus Catalog (https://wwwn.cdc.gov/arbocat/). Where a taxa was identified as an “Arbovirus” by the Arbovirus Catalog it was assigned “yes”, otherwise for those listed as “potential arboviruses”, “probable arboviruses”, or those not present in the catalogue, literature on this taxa was reviewed for evidence of vector association. Three main criteria were considered: (1) whether the virus replicated in both invertebrate and vertebrate cells; (2) the phylogenetic position of the virus, i.e., is the virus in the middle of an insect-specific clade? And (3) consensus among the literature on the possibility of the virus being vectored. The assigned vector status for each taxa and the underlying evidence for this is provided in STable. 3.

#### Evaluation of topological concordance

To determine the most robust NS5b phylogeny, alignments (pre- and post-trimming) (n = 225) were examined for the presence of conical RdRp motifs, misalignments, and overall pairwise identity and length. The resultant tree topology and branch support were examined in FigTree (v1.4.4) ^82^. This analysis was combined with comparisons of genome composition and to previous *Flaviviridae* phylogenies ^7,37,68^ to identify the most concordant topology across the multiple parameters tested. To supplement this, the R package *treespace* (v1.1.4.2) ^83^ was used to conduct a principal component analysis (PCoA) with the goal of identifying clusters of similar trees and assessing whether the selected topology is consistent with the median topology. Accordingly, Kendall–Colijn (KC) distance was calculated for each tree and used for the PCoA, with two principal components retained ^84^. To identify discrete clusters of related trees, pairwise distances were mapped into four clusters using hierarchical clustering (Ward’s method) ^85^. Manual and distance-based inspection revealed that the alignment method drove variation in tree topology and branch lengths between phylogenies. Specifically, tree topologies and their corresponding phylogenetic distances derived from Clustal Omega were frequently topologically discordant compared to those generated by MAFFT and MUSCLE; as such, these phylogenies were excluded and the PCoA was recalculated. Geometric median trees were generated from each cluster and alignment method and used to inform the selection of the final phylogeny. This phylogeny was aligned using MUSCLE with a trimAl consensus and gap threshold value of 5 and 0.9, respectively, and based on the LG+F+R10 amino acid substitution model.

#### RNase T2 phylogeny

To infer the evolutionary history of the RNase T2 protein, sequences were obtained from the GenBank protein database for conserved domains using the queries “taxid 238513” and “taxid 238220”, literature searches ^86,87^, these were supplemented with structurally homologous protein clusters identified using the AlphaFold database FoldSeek clusters server ^88^.

To identify unannotated RNase T2-like sequences in virus genomes, a NCBI web protein BLAST (https://blast.ncbi.nlm.nih.gov/Blast.cgi) was employed with RNase T2 sequences used as a query against the NR clustered database (as of June 2023) ^89^, employing the BLOSUM45 matrix and with taxonomy limited to the group “Viruses” (taxid:10239). The HMM search webserver (v2.41.2) ^90^ was used to identify additional viral T2 RNase-like sequences. An alignment of RNase T2 sequences was used as a query against the Reference Proteomes, UniProtKB, SwissProt and PDB databases (as of June 2023), with results again limited to “Viruses” (taxid:10239). For both methods, if new virus sequences were detected, they were manually inspected for the presence of RNase T2 motifs and, in turn, used as queries. To estimate the RNase T2 phylogeny, non-viral sequences were clustered at 80% amino acid identity using DIAMOND cluster (v.2.0.9) ^91^ with default parameters and aligned with the viral sequences using MAFFT and a maximum likelihood phylogeny as described above.

#### Structure-guided glycoprotein phylogenies

Structurally aligned residues from FoldSeek were used to construct protein multiple sequence alignments that reflect the structural homology between glycoproteins (Fig. 3). However, due to the arbitrary fragmentation of protein sequences into the 300 residue blocks, our query structures represent overlapping truncated segments of the true glycoproteins. To reconstruct the pairwise alignment of the full query glycoprotein matching a reference sequence, we filtered the results by a single reference and retained the five query hits with the highest bitscore. An e-value threshold of 0.01 was also applied to exclude false hits. If this list contained segments that were adjacent in the polyprotein to the hit with the highest bitscore, then the hits were ordered by their block number and their pairwise alignments to the reference were combined. For non-matching overlapping alignment segments, the alignment with the highest bitscore was used. Amino acids in the query proteins that did not match the reference sequence were removed and all hit pairwise alignments, now constrained to the reference sequence’s length, were combined to make single, structure-aware MSAs of the query hits against the given reference(s).

Our reference library contained multiple instances of each glycoprotein target, such that reference structures were selected to capture the greatest number of hits across the *Flaviviridae*. In the case of the E glycoprotein, the DENV serotype 1 AlphaFold structure yielded 251 matches including the majority of hits across the orthoflaviviruses, jingmenviruses and LGF. For E1 and E2, no single reference could match a substantial number of viruses from all three of the *Hepaci*-, *Pegi*- and *Pestivirus* groups. Hence, an additional alignment strategy was implemented for these proteins. We first performed a FoldSeek search using the library of reference structures as both the query and the database set. This provided us with structure-aware pairwise alignments between each of the reference structures. We then used these aligned references to guide the combination of MSAs containing our protein foldome hits against two reference structures, chosen to maximise the coverage of viral species. For E1 we combined hits against the Wenling shark virus (WSV) and BVDV-1 AlphaFold structures (matching 157 hepaci-/pegiviruses and 34 pestiviruses, respectively). For E2, the Thieler’s disease-associated virus (TDAV) and BVDV-1 AlphaFold structures were combined (matching 136 hepaci-/pegiviruses and 33 pestiviruses, respectively).

The resulting structure-aware MSAs were used for inferring ML phylogenies with IQ-TREE (version 2.1.3) ^71^. The substitution model was selected with ModelFinder ^72^ (option ‘-m TEST’) and 10,000 ultrafast bootstrap iterations were implemented for assessing node support. To improve the visualisation, internal nodes with support below 45% were dropped using the ETE3 Python package ^75^. Upon manual inspection of the phylogenies, it was observed that the Dipteran jingmen-related virus (isolate OKIAV332) and the Wuhan aphid virus 1 (strain WHYC-1) E glycoproteins formed a clade placed within the broader LGF clade in the E tree, with a very long internal branch. Based on the high diversity of these two sequences and their inconsistent topology in the tree, indicative of long branch attraction, this clade was manually removed from the final E phylogeny.

## Key resource table

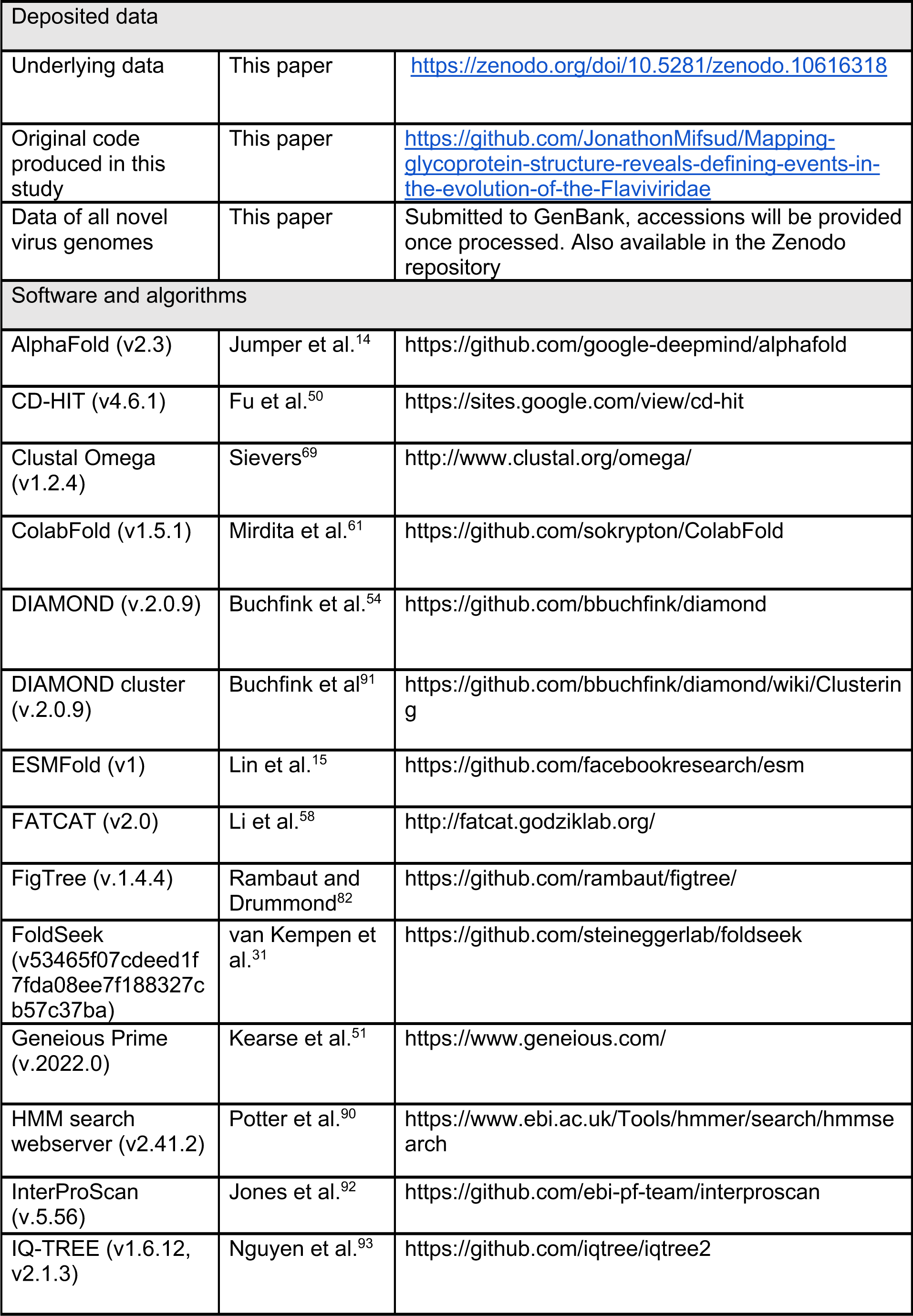
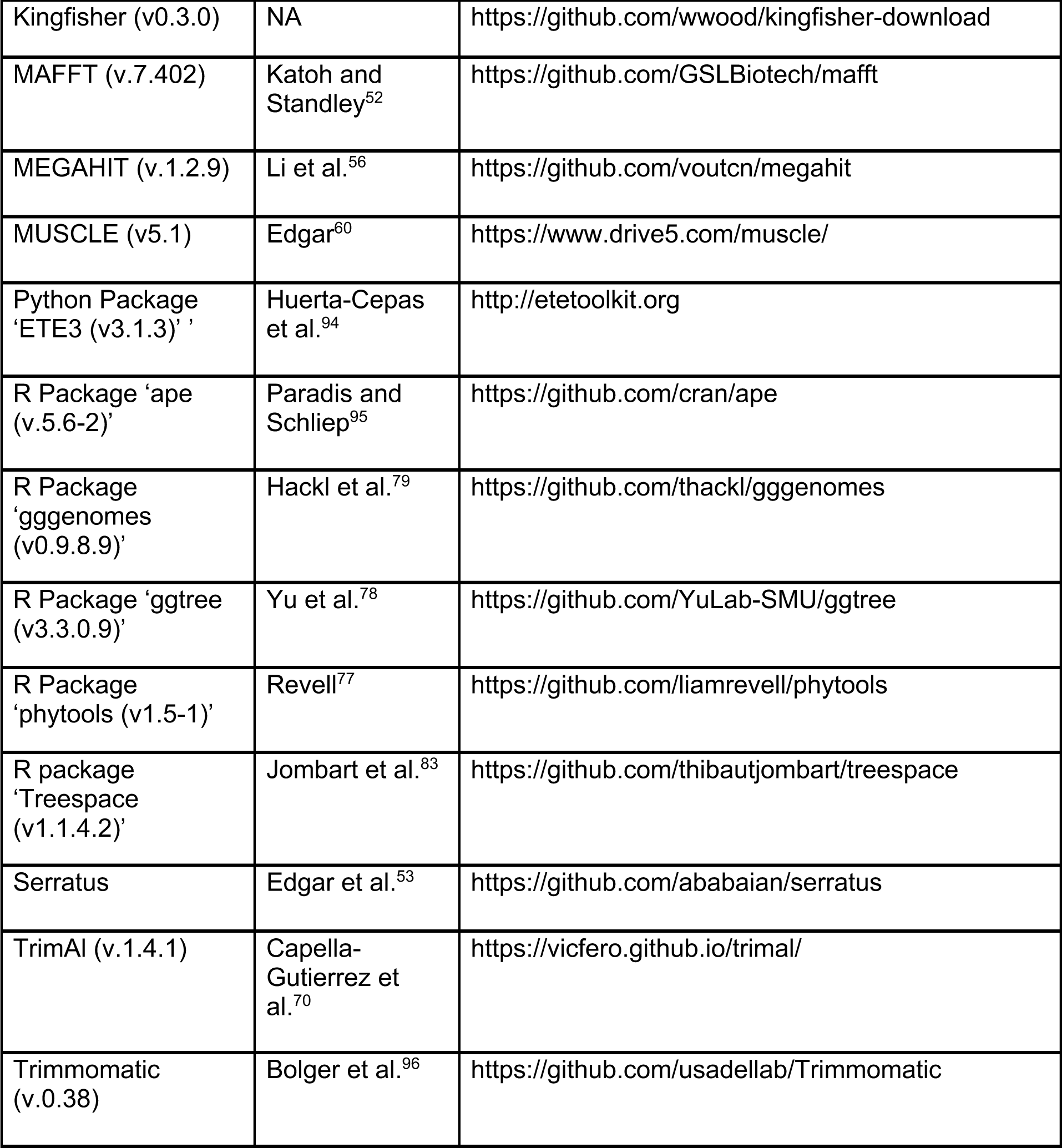

## References

1. Hubálek, Z., and Halouzka, J. (1999). West Nile fever--a reemerging mosquito-borne viral disease in Europe. Emerg. Infect. Dis. 5, 643–650.

2. Wang, Z.-D., Wang, B., Wei, F., Han, S.-Z., Zhang, L., Yang, Z.-T., Yan, Y., Lv, X.-L., Li, L., Wang, S.-C., et al. (2019). A New Segmented Virus Associated with Human Febrile Illness in China. N. Engl. J. Med. 380, 2116–2125.

3. Kartashov, M.Y., Gladysheva, A.V., Shvalov, A.N., Tupota, N.L., Chernikova, A.A., Ternovoi, V.A., and Loktev, V.B. (2023). Novel Flavi-like virus in ixodid ticks and patients in Russia. Ticks Tick Borne Dis. 14, 102101.

4. Postler, T.S., Beer, M., Blitvich, B.J., Bukh, J., de Lamballerie, X., Drexler, J.F., Imrie, A., Kapoor, A., Karganova, G.G., Lemey, P., et al. (2023). Renaming of the genus Flavivirus to Orthoflavivirus and extension of binomial species names within the family Flaviviridae. Arch. Virol. 168, 224.

5. Qin, X.-C., Shi, M., Tian, J.-H., Lin, X.-D., Gao, D.-Y., He, J.-R., Wang, J.-B., Li, C.-X., Kang, Y.-J., Yu, B., et al. (2014). A tick-borne segmented RNA virus contains genome segments derived from unsegmented viral ancestors. Proc. Natl. Acad. Sci. U. S. A. 111, 6744–6749.

6. Ladner, J.T., Wiley, M.R., Beitzel, B., Auguste, A.J., Dupuis, A.P., 2nd, Lindquist, M.E., Sibley, S.D., Kota, K.P., Fetterer, D., Eastwood, G., et al. (2016). A Multicomponent Animal Virus Isolated from Mosquitoes. Cell Host Microbe 20, 357–367.

7. Paraskevopoulou, S., Käfer, S., Zirkel, F., Donath, A., Petersen, M., Liu, S., Zhou, X., Drosten, C., Misof, B., and Junglen, S. (2021). Viromics of extant insect orders unveil the evolution of the flavi-like superfamily. Virus Evolution 7. 10.1093/ve/veab030.

8. Kobayashi, K., Atsumi, G., Iwadate, Y., Tomita, R., Chiba, K.-I., Akasaka, S., Nishihara, M., Takahashi, H., Yamaoka, N., Nishiguchi, M., et al. (2013). Gentian Kobu-sho-associated virus: a tentative, novel double-stranded RNA virus that is relevant to gentian Kobu-sho syndrome. J. Gen. Plant Pathol. 79, 56–63.

9. Debat, H., and Bejerman, N. (2023). Two novel flavi-like viruses shed light on the plant-infecting koshoviruses. Arch. Virol. 168, 184.

10. Petrone, M.E., Grove, J., Mifsud, J.C.O., Parry, R.H., Marzinelli, E.M., and Holmes, E.C. (2024). A 39.8kb flavi-like virus uses a novel strategy for overcoming the RNA virus error threshold. bioRxiv, 2024.01.08.574764. 10.1101/2024.01.08.574764.

11. Ferron, F., Sama, B., Decroly, E., and Canard, B. (2021). The enzymes for genome size increase and maintenance of large (+)RNA viruses. Trends Biochem. Sci. 46, 866–877.

12. Shi, M., Lin, X.-D., Vasilakis, N., Tian, J.-H., Li, C.-X., Chen, L.-J., Eastwood, G., Diao, X.-N., Chen, M.-H., Chen, X., et al. (2016). Divergent Viruses Discovered in Arthropods and Vertebrates Revise the Evolutionary History of the Flaviviridae and Related Viruses. J. Virol. 90, 659–669.

13. Garry, C.E., and Garry, R.F. (2020). Proteomics Computational Analyses Suggest That the Envelope Glycoproteins of Segmented Jingmen Flavi-Like Viruses are Class II Viral Fusion Proteins (b-Penetrenes) with Mucin-Like Domains. Viruses 12. 10.3390/v12030260.

14. Jumper, J., Evans, R., Pritzel, A., Green, T., Figurnov, M., Ronneberger, O., Tunyasuvunakool, K., Bates, R., Žídek, A., Potapenko, A., et al. (2021). Highly accurate protein structure prediction with AlphaFold. Nature 596, 583–589.

15. Lin, Z., Akin, H., Rao, R., Hie, B., Zhu, Z., Lu, W., Smetanin, N., Verkuil, R., Kabeli, O., Shmueli, Y., et al. (2023). Evolutionary-scale prediction of atomic-level protein structure with a language model. Science 379, 1123–1130.

16. Kielian, M., and Rey, F.A. (2006). Virus membrane-fusion proteins: more than one way to make a hairpin. Nat. Rev. Microbiol. 4, 67–76.

17. Rey, F.A., Heinz, F.X., Mandl, C., Kunz, C., and Harrison, S.C. (1995). The envelope glycoprotein from tick-borne encephalitis virus at 2 A resolution. Nature 375, 291–298.

18. Dessau, M., and Modis, Y. (2013). Crystal structure of glycoprotein C from Rift Valley fever virus. Proc. Natl. Acad. Sci. U. S. A. 110, 1696–1701.

19. Fédry, J., Liu, Y., Péhau-Arnaudet, G., Pei, J., Li, W., Tortorici, M.A., Traincard, F., Meola, A., Bricogne, G., Grishin, N.V., et al. (2017). The Ancient Gamete Fusogen HAP2 Is a Eukaryotic Class II Fusion Protein. Cell 168, 904–915.e10.

20. Guardado-Calvo, P., and Rey, F.A. (2021). The Viral Class II Membrane Fusion Machinery: Divergent Evolution from an Ancestral Heterodimer. Viruses 13. 10.3390/v13122368.

21. Li, L., Lok, S.-M., Yu, I.-M., Zhang, Y., Kuhn, R.J., Chen, J., and Rossmann, M.G. (2008). The flavivirus precursor membrane-envelope protein complex: structure and maturation. Science 319, 1830–1834.

22. Vaney, M.-C., Dellarole, M., Duquerroy, S., Medits, I., Tsouchnikas, G., Rouvinski, A., England, P., Stiasny, K., Heinz, F.X., and Rey, F.A. (2022). Evolution and activation mechanism of the flavivirus class II membrane-fusion machinery. Nat. Commun. 13, 3718.

23. El Omari, K., Iourin, O., Harlos, K., Grimes, J.M., and Stuart, D.I. (2013). Structure of a pestivirus envelope glycoprotein E2 clarifies its role in cell entry. Cell Rep. 3, 30–35.

24. Li, Y., Wang, J., Kanai, R., and Modis, Y. (2013). Crystal structure of glycoprotein E2 from bovine viral diarrhea virus. Proc. Natl. Acad. Sci. U. S. A. 110, 6805–6810.

25. Kong, L., Giang, E., Nieusma, T., Kadam, R.U., Cogburn, K.E., Hua, Y., Dai, X., Stanfield, R.L., Burton, D.R., Ward, A.B., et al. (2013). Hepatitis C virus E2 envelope glycoprotein core structure. Science 342, 1090–1094.

26. Khan, A.G., Whidby, J., Miller, M.T., Scarborough, H., Zatorski, A.V., Cygan, A., Price, A.A., Yost, S.A., Bohannon, C.D., Jacob, J., et al. (2014). Structure of the core ectodomain of the hepatitis C virus envelope glycoprotein 2. Nature 509, 381–384.

27. Aitkenhead, H., Riedel, C., Cowieson, N., Rümenapf, H.T., Stuart, D.I., and El Omari, K. (2023). Structural comparison of typical and atypical E2 pestivirus glycoproteins. Structure. 10.1016/j.str.2023.12.003.

28. Torrents de la Peña, A., Sliepen, K., Eshun-Wilson, L., Newby, M.L., Allen, J.D., Zon, I., Koekkoek, S., Chumbe, A., Crispin, M., Schinkel, J., et al. (2022). Structure of the hepatitis C virus E1E2 glycoprotein complex. Science 378, 263–269.

29. Metcalf, M.C., Janus, B.M., Yin, R., Wang, R., Guest, J.D., Pozharski, E., Law, M., Mariuzza, R.A., Toth, E.A., Pierce, B.G., et al. (2023). Structure of engineered hepatitis C virus E1E2 ectodomain in complex with neutralizing antibodies. Nat. Commun. 14, 3980.

30. Oliver, M.R., Toon, K., Lewis, C.B., Devlin, S., Gifford, R.J., and Grove, J. (2023). Structures of the Hepaci-, Pegi-, and Pestiviruses envelope proteins suggest a novel membrane fusion mechanism. PLoS Biol. 21, e3002174.

31. van Kempen, M., Kim, S.S., Tumescheit, C., Mirdita, M., Lee, J., Gilchrist, C.L.M., Söding, J., and Steinegger, M. (2023). Fast and accurate protein structure search with Foldseek. Nat. Biotechnol., 1–4.

32. Urayama, S.-I., Takaki, Y., and Nunoura, T. (2016). FLDS: A Comprehensive dsRNA Sequencing Method for Intracellular RNA Virus Surveillance. Microbes Environ. 31, 33–40.

33. Hou, X., He, Y., Fang, P., Mei, S.-Q., Xu, Z., Wu, W.-C., Tian, J.-H., Zhang, S., Zeng, Z.-Y., Gou, Q.-Y., et al. (2023). Artificial intelligence redefines RNA virus discovery. bioRxiv, 2023.04.18.537342.

34. Chen, Y.-M., Sadiq, S., Tian, J.-H., Chen, X., Lin, X.-D., Shen, J.-J., Chen, H., Hao, Z.-Y., Wille, M., Zhou, Z.-C., et al. (2022). RNA viromes from terrestrial sites across China expand environmental viral diversity. Nat Microbiol 7, 1312–1323.

35. Modis, Y., Ogata, S., Clements, D., and Harrison, S.C. (2004). Structure of the dengue virus envelope protein after membrane fusion. Nature 427, 313–319.

36. Krey, T., Bontems, F., Vonrhein, C., Vaney, M.-C., Bricogne, G., Rümenapf, T., and Rey, F.A. (2012). Crystal structure of the pestivirus envelope glycoprotein E(rns) and mechanistic analysis of its ribonuclease activity. Structure 20, 862–873.

37. Bamford, C.G.G., de Souza, W.M., Parry, R., and Gifford, R.J. (2022). Comparative analysis of genome-encoded viral sequences reveals the evolutionary history of flavivirids (family Flaviviridae). Virus Evolution 8. 10.1093/ve/veac085.

38. Mushegian, A. (2022). Methyltransferases of Riboviria. Biomolecules 12. 10.3390/biom12091247.

39. Ding, Q., von Schaewen, M., and Ploss, A. (2014). The impact of hepatitis C virus entry on viral tropism. Cell Host Microbe 16, 562–568.

40. Pfaender, S., Cavalleri, J.M.V., Walter, S., Doerrbecker, J., Campana, B., Brown, R.J.P., Burbelo, P.D., Postel, A., Hahn, K., Anggakusuma, et al. (2015). Clinical course of infection and viral tissue tropism of hepatitis C virus-like nonprimate hepaciviruses in horses. Hepatology 61, 447–459.

41. Costa, V.A., Ronco, F., Mifsud, J.C.O., and Harvey, E. (2023). Host adaptive radiation is associated with rapid virus diversification and cross-species transmission in African cichlid fishes. bioRxiv.

42. Mifsud, J.C.O., Costa, V.A., Petrone, M.E., Marzinelli, E.M., Holmes, E.C., and Harvey, E. (2022). Transcriptome mining extends the host range of the Flaviviridae to non-bilaterians. Virus Evolution 9. 10.1093/ve/veac124.

43. Balsitis, S.J., Coloma, J., Castro, G., Alava, A., Flores, D., McKerrow, J.H., Beatty, P.R., and Harris, E. (2009). Tropism of dengue virus in mice and humans defined by viral nonstructural protein 3-specific immunostaining. Am. J. Trop. Med. Hyg. 80, 416–424.

44. Kong, Y., Zhang, G., Jiang, L., Wang, P., Zhang, S., Zheng, X., and Li, Y. (2022). Metatranscriptomics Reveals the Diversity of the Tick Virome in Northwest China. Microbiol Spectr 10, e0111522.

45. Costa, V.A., Bellwood, D.R., Mifsud, J.C.O., Geoghegan, J.L., Holmes, E.C., and Harvey, E. (2022). Limited Cross-Species Virus Transmission in a Spatially Restricted Coral Reef Fish Community. bioRxiv, 2022.05.17.492384.

46. Perveen, N., Kundu, B., Sudalaimuthuasari, N., Al-Maskari, R.S., Muzaffar, S.B., and Al-Deeb, M.A. (2023). Virome diversity of Hyalomma dromedarii ticks collected from camels in the United Arab Emirates. Vet World 16, 439–448.

47. Guo, G., Wang, M., Zhou, D., He, X., Han, P., Chen, G., Zeng, J., Liu, Z., Wu, Y., Weng, S., et al. (2023). Virome Analysis Provides an Insight into the Viral Community of Chinese Mitten Crab Eriocheir sinensis. Microbiol Spectr 11, e0143923.

48. Dunay, E., Owens, L.A., Dunn, C.D., Rukundo, J., Atencia, R., Cole, M.F., Cantwell, A., Emery Thompson, M., Rosati, A.G., and Goldberg, T.L. (2023). Viruses in sanctuary chimpanzees across Africa. Am. J. Primatol. 85, e23452.

49. Elbadry, M.A., Efstathion, C.A., Qualls, W.A., Tagliamonte, M.S., Alam, M.M., Khan, M.S.R., Ryan, S.J., Xue, R.D., Charrel, R.N., Bangonan, L., et al. (2023). Diversity and Genetic Reassortment of Keystone Virus in Mosquito Populations in Florida. Am. J. Trop. Med. Hyg. 108, 1256–1263.

50. Fu, L., Niu, B., Zhu, Z., Wu, S., and Li, W. (2012). CD-HIT: accelerated for clustering the next-generation sequencing data. Bioinformatics 28, 3150–3152.

51. Kearse, M., Moir, R., Wilson, A., Stones-Havas, S., Cheung, M., Sturrock, S., Buxton, S., Cooper, A., Markowitz, S., and Duran, C. (2012). Geneious Basic: an integrated and extendable desktop software platform for the organization and analysis of sequence data. Bioinformatics 28, 1647–1649.

52. Katoh, K., and Standley, D.M. (2013). MAFFT multiple sequence alignment software version 7: improvements in performance and usability. Mol. Biol. Evol. 30, 772–780.

53. Edgar, R.C., Taylor, J., Lin, V., Altman, T., Barbera, P., Meleshko, D., Lohr, D., Novakovsky, G., Buchfink, B., and Al-Shayeb, B. (2022). Petabase-scale sequence alignment catalyses viral discovery. Nature 602, 142–147.

54. Buchfink, B., Reuter, K., and Drost, H.-G. (2021). Sensitive protein alignments at tree-of-life scale using DIAMOND. Nat. Methods 18, 366–368.

55. Mifsud, J.C.O. (2023). BatchArtemisSRAMiner: v1.0.0. 10.5281/ZENODO.8417951.

56. Li, D., Liu, C.-M., Luo, R., Sadakane, K., and Lam, T.-W. (2015). MEGAHIT: an ultra-fast single-node solution for large and complex metagenomics assembly via succinct de Bruijn graph. Bioinformatics 31, 1674–1676.

57. Meng, E.C., Goddard, T.D., Pettersen, E.F., Couch, G.S., Pearson, Z.J., Morris, J.H., and Ferrin, T.E. (2023). UCSF ChimeraX: Tools for structure building and analysis. Protein Sci. 32, e4792.

58. Li, Z., Jaroszewski, L., Iyer, M., Sedova, M., and Godzik, A. (2020). FATCAT 2.0: towards a better understanding of the structural diversity of proteins. Nucleic Acids Res. 48, W60–W64.

59. Teufel, F., Almagro Armenteros, J.J., Johansen, A.R., Gíslason, M.H., Pihl, S.I., Tsirigos, K.D., Winther, O., Brunak, S., von Heijne, G., and Nielsen, H. (2022). SignalP 6.0 predicts all five types of signal peptides using protein language models. Nat. Biotechnol. 40, 1023–1025.

60. Edgar, R.C. (2022). Muscle5: High-accuracy alignment ensembles enable unbiased assessments of sequence homology and phylogeny. Nat. Commun. 13, 6968.

61. Mirdita, M., Schütze, K., Moriwaki, Y., Heo, L., Ovchinnikov, S., and Steinegger, M. (2022). ColabFold: making protein folding accessible to all. Nat. Methods 19, 679–682.

62. Letunic, I., and Bork, P. (2021). Interactive Tree Of Life (iTOL) v5: an online tool for phylogenetic tree display and annotation. Nucleic Acids Res. 49, W293–W296.

63. Renner, M., Dejnirattisai, W., Carrique, L., Martin, I.S., Karia, D., Ilca, S.L., Ho, S.F., Kotecha, A., Keown, J.R., Mongkolsapaya, J., et al. (2021). Flavivirus maturation leads to the formation of an occupied lipid pocket in the surface glycoproteins. Nat. Commun. 12, 1238.

64. Egloff, M.-P., Benarroch, D., Selisko, B., Romette, J.-L., and Canard, B. (2002). An RNA cap (nucleoside-2’-O-)-methyltransferase in the flavivirus RNA polymerase NS5: crystal structure and functional characterization. EMBO J. 21, 2757–2768.

65. Noble, C.G., Lim, S.P., Arora, R., Yokokawa, F., Nilar, S., Seh, C.C., Wright, S.K., Benson, T.E., Smith, P.W., and Shi, P.-Y. (2016). A Conserved Pocket in the Dengue Virus Polymerase Identified through Fragment-based Screening. J. Biol. Chem. 291, 8541–8548.

66. Jia, H., Zhong, Y., Peng, C., and Gong, P. (2022). Crystal Structures of Flavivirus NS5 Guanylyltransferase Reveal a GMP-Arginine Adduct. J. Virol. 96, e0041822.

67. Walls, A.C., Park, Y.-J., Tortorici, M.A., Wall, A., McGuire, A.T., and Veesler, D. (2020). Structure, Function, and Antigenicity of the SARS-CoV-2 Spike Glycoprotein. Cell 181, 281–292.e6.

68. Dong, X., Wang, G., Hu, T., Li, J., Li, C., Cao, Z., Shi, M., Wang, Y., Zou, P., Song, J., et al. (2021). A Novel Virus of Flaviviridae Associated with Sexual Precocity in Macrobrachium rosenbergii. mSystems 6, e00003–e00021.

69. Sievers, F., Wilm, A., Dineen, D., Gibson, T.J., Karplus, K., Li, W., Lopez, R., McWilliam, H., Remmert, M., and Söding, J. (2011). Fast, scalable generation of high-quality protein multiple sequence alignments using Clustal Omega. Mol. Syst. Biol. 7, 539.

70. Capella-Gutiérrez, S., Silla-Martínez, J.M., and Gabaldón, T. (2009). trimAl: a tool for automated alignment trimming in large-scale phylogenetic analyses. Bioinformatics 25, 1972–1973.

71. Minh, B.Q., Schmidt, H.A., Chernomor, O., Schrempf, D., Woodhams, M.D., Von Haeseler, A., and Lanfear, R. (2020). IQ-TREE 2: new models and efficient methods for phylogenetic inference in the genomic era. Mol. Biol. Evol. 37, 1530–1534.

72. Kalyaanamoorthy, S., Minh, B.Q., Wong, T.K.F., Von Haeseler, A., and Jermiin, L.S. (2017). ModelFinder: fast model selection for accurate phylogenetic estimates. Nat. Methods 14, 587–589.

73. Le, T.K., and Vinh, L.S. (2020). FLAVI: an amino acid substitution model for flaviviruses. J. Mol. Evol. 88, 445–452.

74. Guindon, S., Dufayard, J.F., Lefort, V., Anisimova, M., Hordijk, W., and Gascuel, O. (2010). New Algorithms and Methods to Estimate Maximum-Likelihood Phylogenies: Assessing the Performance of PhyML 3.0. Syst. Biol. 59, 307–321.

75. Hoang, D.T., Chernomor, O., von Haeseler, A., Minh, B.Q., and Vinh, L.S. (2017). UFBoot2: Improving the Ultrafast Bootstrap Approximation. Mol. Biol. Evol. 35, 518–522.

76. Simmonds, P., Becher, P., Bukh, J., Gould, E.A., Meyers, G., Monath, T., Muerhoff, S., Pletnev, A., Rico-Hesse, R., and Smith, D.B. (2017). ICTV virus taxonomy profile: Flaviviridae. J. Gen. Virol. 98, 2.

77. Revell, L.J. (2023). phytools 2.0: An updated R ecosystem for phylogenetic comparative methods (and other things). bioRxiv, 2023.03. 08.531791.

78. Yu, G., Smith, D.K., Zhu, H., Guan, Y., and Lam, T.T. (2017). ggtree: an R package for visualization and annotation of phylogenetic trees with their covariates and other associated data. Methods Ecol. Evol. 8, 28–36.

79. Thomas Hackl, M.J.A. (2022). gggenomes: A Grammar of Graphics for Comparative Genomics.

80. Winter, D.J. (2017). rentrez: An R package for the NCBI eUtils API (PeerJ Preprints).

81. Chamberlain, S.A., and Szöcs, E. (2013). taxize: taxonomic search and retrieval in R. F1000Res. 2, 191.

82. Rambaut, A., and Drummond, A.J. (2012). FigTree: Tree figure drawing tool, version 1.4.0.

83. Jombart, T., Kendall, M., Almagro-Garcia, J., and Colijn, C. (2017). treespace: Statistical exploration of landscapes of phylogenetic trees. Mol. Ecol. Resour. 17, 1385–1392.

84. Kendall, M., and Colijn, C. (2016). Mapping phylogenetic trees to reveal distinct patterns of evolution. Mol. Biol. Evol. 33, 2735–2743.

85. Legendre, P., and Legendre, L. (2012). Numerical ecology (Elsevier).

86. Saberi, A., Gulyaeva, A.A., Brubacher, J.L., Newmark, P.A., and Gorbalenya, A.E. (2018). A planarian nidovirus expands the limits of RNA genome size. PLoS Pathog. 14, e1007314.

87. Rolland, C., La Scola, B., and Levasseur, A. (2020). How Tupanvirus Degrades the Ribosomal RNA of Its Amoebal Host? The Ribonuclease T2 Track. Front. Microbiol. 11, 1691.

88. Barrio-Hernandez, I., Yeo, J., Jänes, J., Mirdita, M., Gilchrist, C.L.M., Wein, T., Varadi, M., Velankar, S., Beltrao, P., and Steinegger, M. (2023). Clustering predicted structures at the scale of the known protein universe. Nature 622, 637–645.

89. Steinegger, M., and Söding, J. (2017). MMseqs2 enables sensitive protein sequence searching for the analysis of massive data sets. Nat. Biotechnol. 35, 1026–1028.

90. Potter, S.C., Luciani, A., Eddy, S.R., Park, Y., Lopez, R., and Finn, R.D. (2018). HMMER web server: 2018 update. Nucleic Acids Res. 46, W200–W204.

91. Buchfink, B., Ashkenazy, H., Reuter, K., Kennedy, J.A., and Drost, H.-G. (2023). Sensitive clustering of protein sequences at tree-of-life scale using DIAMOND DeepClust. bioRxiv, 2023.01.24.525373.

92. Jones, P., Binns, D., Chang, H.-Y., Fraser, M., Li, W., McAnulla, C., McWilliam, H., Maslen, J., Mitchell, A., and Nuka, G. (2014). InterProScan 5: genome-scale protein function classification. Bioinformatics 30, 1236–1240.

93. Nguyen, L.-T., Schmidt, H.A., Von Haeseler, A., and Minh, B.Q. (2015). IQ-TREE: a fast and effective stochastic algorithm for estimating maximum-likelihood phylogenies. Mol. Biol. Evol. 32, 268–274.

94. Huerta-Cepas, J., Serra, F., and Bork, P. (2016). ETE 3: Reconstruction, Analysis, and Visualization of Phylogenomic Data. Mol. Biol. Evol. 33, 1635–1638.

95. Paradis, E., and Schliep, K. (2019). ape 5.0: an environment for modern phylogenetics and evolutionary analyses in R. Bioinformatics 35, 526–528.

96. Bolger, A.M., Lohse, M., and Usadel, B. (2014). Trimmomatic: a flexible trimmer for Illumina sequence data. Bioinformatics 30, 2114–2120.

