## Supplementary Materials for "Mapping glycoprotein structure reveals defining events in the evolution of the *Flaviviridae*"

### Supplementary Figure Legends

**Supplementary Figure 1. Two-dimensional MDS plot of the NS5b phylogeny variations A.** Coloured by alignment method **B.** Excluding trees generated from Clustal Omega alignments. Points, which represent individual phylogenies, are colour-coded based on the clusters identified using the 'findGroves' function. Point shapes indicate the alignment software, the trimAl gap and consensus thresholds, and the substitution model applied. An arrow signifies the master phylogeny chosen for further analysis.

**Supplementary Figure 2. NS5b consensus phylogeny (Tree 18) with tip labels.** The complete NS5b phylogeny (underlying Fig. 1A & 2A) split by the major *Flaviviridae* lineages **A.** *Orthoflavivirus/Jingmenvirus*, **B.** Large genome flavivirus/*Pestivirus* and **C.** *Hepacivirus/Pegivirus*. A scale bar denoting the number of amino acid substitutions per site. Node support (SH-aLRT >=80% & UFboot >=95%) is indicated by a black circle while intermediate support (SH-aLRT <80% & UFboot >=95% or SH-aLRT >80% & UFboot <95%) is indicated by a half filled circle corresponding to the indice above the support threshold.

**Supplementary Figure 3. Analysis of environmental pesti-like viruses. A.** Large genome flavivirus/*Pestivirus* subset of the *Flaviviridae* phylogeny (Tree 18) collapsed to highlight the environmental pesti-like viruses for which no glycoproteins were identified. A scale bar denoting the number of amino acid substitutions per site. **B.** Genome organisation is provided for each species, with annotations based on conserved domain sequence searches. **C.** FoldSeek e-value heatmaps for the indicated reference proteins, values are log transformed and colourcolor-coded as shown in the key. For E1, E2, prM and E, the values represent summary e-values after comparison with a range of relevant reference structures, as described in the methods.

**Supplementary Table 1.** *Flaviviridae* sequence metadata including clade designations and GenBank nucleotide accession numbers (attached as a .csv).

**Supplementary Table 2.** Combination of sequence alignment, quality trimming methods, and amino acid substitution models used to infer the NS5b phylogenies (attached as a .csv).

**Supplementary Table 3.** *Flaviviridae* host association and vector status metadata related to Fig. 2C (attached as a .csv).

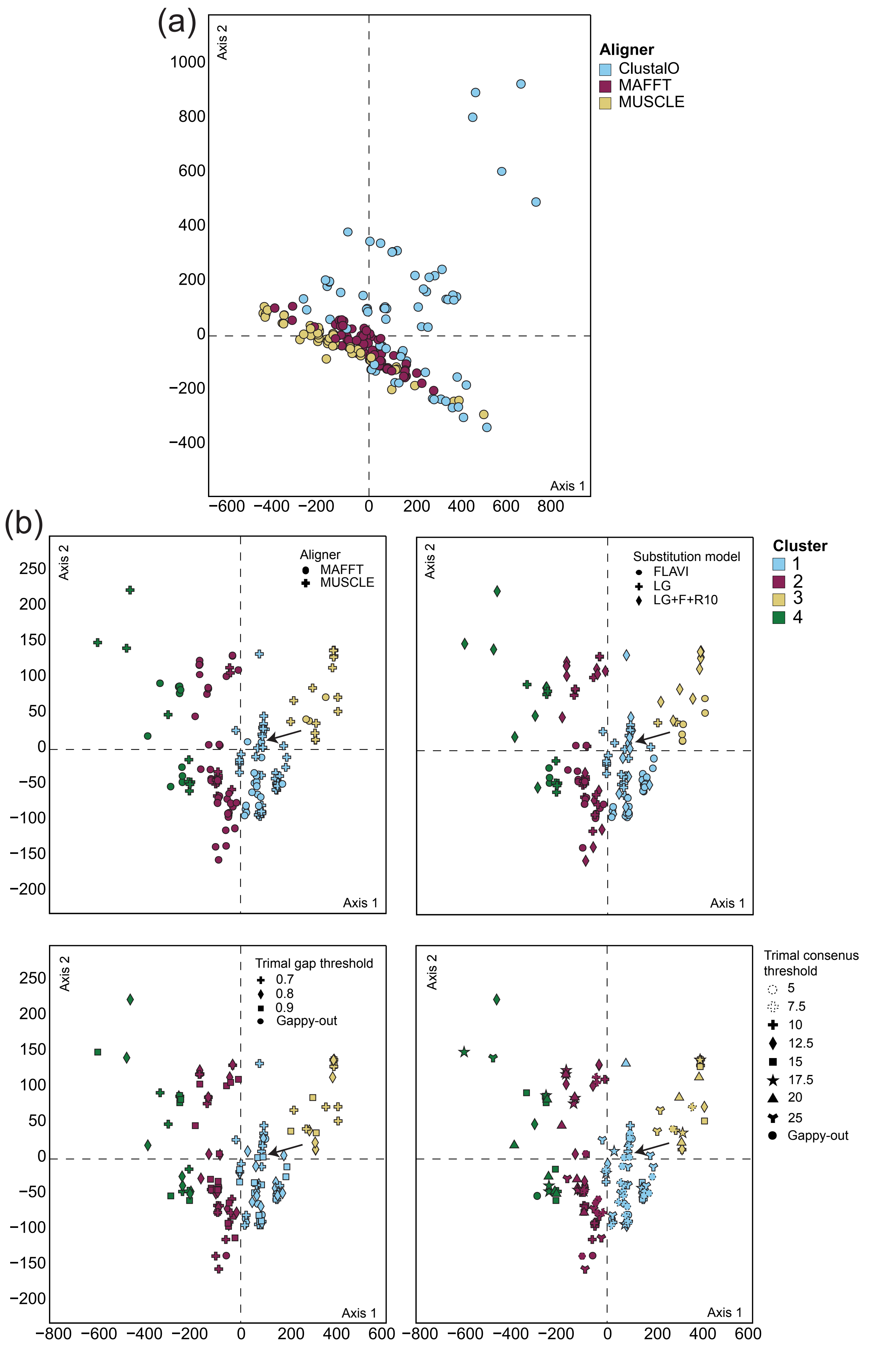

(a)
Axis 2
Axis 1
1000
800
600
400
200
0
−200
−400
−600 −400 −200 0 200 400 600 800
Aligner
ClustalO
MAFFT
MUSCLE
(b)
Aligner
MAFFT
MUSCLE
Substitution model
FLAVI
LG
LG+F+R10
Cluster
1
2
3
4
Trimal gap threshold
0.7
0.8
0.9
Gappy-out
Trimal consenus threshold
5
7.5
10
12.5
15
17.5
20
25
Gappy-out
250
200
150
100
50
0
−50
−100
−150
−200
−800 −600 −400 −200 0 200 400 600


Figure S1

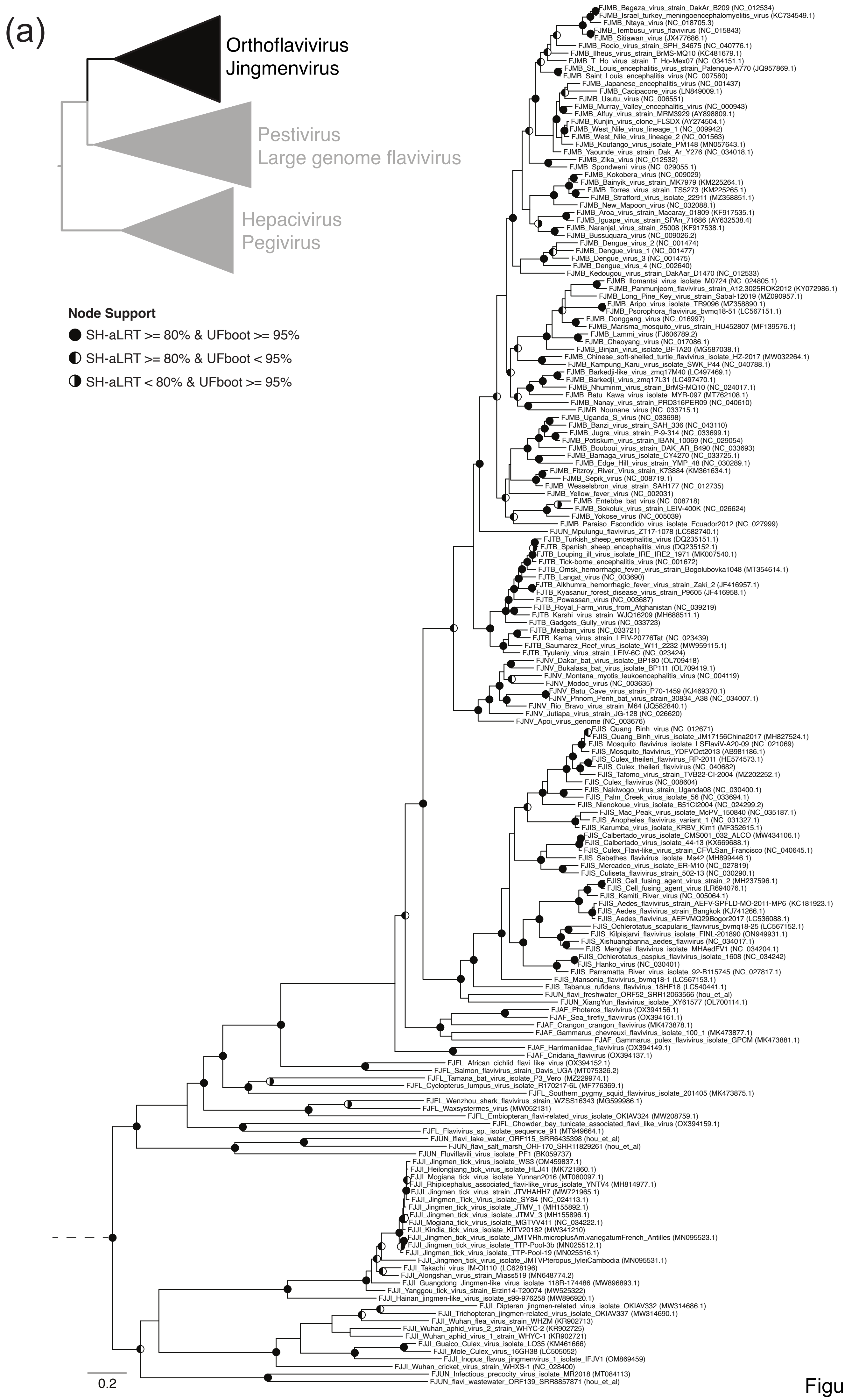

(a)
Orthoflavivirus
Jingmenvirus
Pestivirus
Large genome flavivirus
Hepacivirus
Pegivirus
Node Support
SH-aLRT >= 80% & UFboot >= 95%
SH-aLRT >= 80% & UFboot < 95%
SH-aLRT < 80% & UFboot >= 95%
FJMB_Bagaza_virus_strain_DakAr_B209 (NC_012534)
FJMB_Israel_turkey_meningoencephalomyelitis_virus (KC734549.1)
FJMB_Ntaya_virus (NC_018705.3)
FJMB_Tembusu_virus_flavivirus (NC_015843)
FJMB_Sitiawan_virus (JX477686.1)
FJMB_Rocio_virus_strain_SPH_34675 (NC_040776.1)
FJMB_Ilheus_virus_strain_BrMS-MQ10 (KC481679.1)
FJMB_T_Ho_virus_strain_T_Ho-Mex07 (NC_034151.1)
FJMB_St._Louis_encephalitis_virus_strain_Palenque-A770 (JQ957869.1)
FJMB_Saint_Louis_encephalitis_virus (NC_007580)
FJMB_Japanese_encephalitis_virus (NC_001437)
FJMB_Cacipacore_virus (LN849009.1)
FJMB_Usutu_virus (NC_006551)
FJMB_Murray_Valley_encephalitis_virus (NC_000943)
FJMB_Alfuy_virus_strain_MRM3929 (AY898809.1)
FJMB_Kunjin_virus_clone_FLSDX (AY274504.1)
FJMB_West_Nile_virus_lineage_1 (NC_009942)
FJMB_West_Nile_virus_lineage_2 (NC_001563)
FJMB_Koutango_virus_isolate_PM148 (MN057643.1)
FJMB_Yaounde_virus_strain_Dak_Ar_Y276 (NC_034018.1)
FJMB_Zika_virus (NC_012532)
FJMB_Spondweni_virus (NC_029055.1)
FJMB_Kokobera_virus (NC_009029)
FJMB_Bainyik_virus_strain_MK7979 (KM225264.1)
FJMB_Torres_virus_strain_TS5273 (KM225265.1)
FJMB_Stratford_virus_isolate_22911 (MZ358851.1)
FJMB_New_Mapoon_virus (NC_032088.1)
FJMB_Aroa_virus_strain_Macaray_01809 (KF917535.1)
FJMB_Iguape_virus_strain_SPAn_71686 (AY632538.4)
FJMB_Naranjal_virus_strain_25008 (KF917538.1)
FJMB_Bussuquara_virus (NC_009026.2)
FJMB_Dengue_virus_2 (NC_001474)
FJMB_Dengue_virus_1 (NC_001477)
FJMB_Dengue_virus_3 (NC_001475)
FJMB_Dengue_virus_4 (NC_002640)
FJMB_Kedougou_virus_strain_DakAar_D1470 (NC_012533)
FJMB_Ilomantsi_virus_isolate_M0724 (NC_024805.1)
FJMB_Panmunjeom_flavivirus_strain_A12.3025ROK2012 (KY072986.1)
FJMB_Long_Pine_Key_virus_strain_Sabal-12019 (MZ090957.1)
FJMB_Aripo_virus_isolate_TR9096 (MZ358890.1)
FJMB_Psorophora_flavivirus_bvmq18-51 (LC567151.1)
FJMB_Donggang_virus (NC_016997)
FJMB_Marisma_mosquito_virus_strain_HU452807 (MF139576.1)
FJMB_Lammi_virus (FJ606789.2)
FJMB_Chaoyang_virus (NC_017086.1)
FJMB_Binjari_virus_isolate_BFTA20 (MG587038.1)
FJMB_Chinese_soft-shelled_turtle_flavivirus_isolate_HZ-2017 (MW032264.1)
FJMB_Kampung_Karu_virus_isolate_SWK_P44 (NC_040788.1)
FJMB_Barkedji-like_virus_zmq17M40 (LC497469.1)
FJMB_Barkedji_virus_zmq17L31 (LC497470.1)
FJMB_Nhumirim_virus_strain_BrMS-MQ10 (NC_024017.1)
FJMB_Batu_Kawa_virus_isolate_MYR-097 (MT762108.1)
FJMB_Nanay_virus_strain_PRD316PER09 (NC_040610)
FJMB_Nounane_virus (NC_033715.1)
FJMB_Uganda_S_virus (NC_033698)
FJMB_Banzi_virus_strain_SAH_336 (NC_043110)
FJMB_Jugra_virus_strain_P-9-314 (NC_033699.1)
FJMB_Potiskum_virus_strain_IBAN_10069 (NC_029054)
FJMB_Bouboui_virus_strain_DAK_AR_B490 (NC_033693)
FJMB_Bamaga_virus_isolate_CY4270 (NC_033725.1)
FJMB_Edge_Hill_virus_strain_YMP_48 (NC_030289.1)
FJMB_Fitzroy_River_Virus_strain_K73884 (KM361634.1)
FJMB_Sepik_virus (NC_008719.1)
FJMB_Wesselsbron_virus_strain_SAH177 (NC_012735)
FJMB_Yellow_fever_virus (NC_002031)
FJMB_Entebbe_bat_virus (NC_008718)
FJMB_Sokoluk_virus_strain_LEIV-400K (NC_026624)
FJMB_Yokose_virus (NC_005039)
FJMB_Paraiso_Escondido_virus_isolate_Ecuador2012 (NC_027999)
FJUN_Mpulungu_flavivirus_ZT17-1078 (LC582740.1)
FJTB_Turkish_sheep_encephalitis_virus (DQ235151.1)
FJTB_Spanish_sheep_encephalitis_virus (DQ235152.1)
FJTB_Louping_ill_virus_isolate_IRE_IRE2_1971 (MK007540.1)
FJTB_Tick-borne_encephalitis_virus (NC_001672)
FJTB_Omsk_hemorrhagic_fever_virus_strain_Bogolubovka1048 (MT354614.1)
FJTB_Langat_virus (NC_003690)
FJTB_Alkhumra_hemorrhagic_fever_virus_strain_Zaki_2 (JF416957.1)
FJTB_Kyasanur_forest_disease_virus_strain_P9605 (JF416958.1)
FJTB_Powassan_virus (NC_003687)
FJTB_Royal_Farm_virus_from_Afghanistan (NC_039219)
FJTB_Karshi_virus_strain_WJQ16209 (MH688511.1)
FJTB_Gadgets_Gully_virus (NC_033723)
FJTB_Meaban_virus (NC_033721)
FJTB_Kama_virus_strain_LEIV-20776Tat (NC_023439)
FJTB_Saumarez_Reef_virus_isolate_W11_2232 (MW959115.1)
FJTB_Tyuleniy_virus_strain_LEIV-6C (NC_023424)
FJNV_Dakar_bat_virus_isolate_BP180 (OL709418)
FJNV_Bukalasa_bat_virus_isolate_BP111 (OL709419.1)
FJNV_Montana_myotis_leukoencephalitis_virus (NC_004119)
FJNV_Modoc_virus (NC_003635)
FJNV_Batu_Cave_virus_strain_P70-1459 (KJ469370.1)
FJNV_Phnom_Penh_bat_virus_strain_30834_A38 (NC_034007.1)
FJNV_Rio_Bravo_virus_strain_M64 (JQ582840.1)
FJNV_Jutiapa_virus_strain_JG-128 (NC_026620)
FJNV_Apoi_virus_genome (NC_003676)
FJIS_Quang_Binh_virus (NC_012671)
FJIS_Quang_Binh_virus_isolate_JM17156China2017 (MH827524.1)
FJIS_Mosquito_flavivirus_isolate_LSFlaviV-A20-09 (NC_021069)
FJIS_Mosquito_flavivirus_YDFVOct2013 (AB981186.1)
FJIS_Culex_theileri_flavivirus_RP-2011 (HE574573.1)
FJIS_Culex_theileri_flavivirus (NC_040682)
FJIS_Tafomo_virus_strain_TVB22-CI-2004 (MZ202252.1)
FJIS_Culex_flavivirus (NC_008604)
FJIS_Nakiwogo_virus_strain_Uganda08 (NC_030400.1)
FJIS_Palm_Creek_virus_isolate_56 (NC_033694.1)
FJIS_Nienokoue_virus_isolate_B51CI2004 (NC_024299.2)
FJIS_Mac_Peak_virus_isolate_McPV_150840 (NC_035187.1)
FJIS_Anopheles_flavivirus_variant_1 (NC_031327.1)
FJIS_Karumba_virus_isolate_KRBV_Kim1 (MF352615.1)
FJIS_Calbertado_virus_isolate_CMS001_032_ALCO (MW434106.1)
FJIS_Calbertado_virus_isolate_44-13 (KX669688.1)
FJIS_Culex_Flavi-like_virus_strain_CFVLSan_Francisco (NC_040645.1)
FJIS_Sabethes_flavivirus_isolate_Ms42 (MH899446.1)
FJIS_Mercadeo_virus_isolate_ER-M10 (NC_027819)
FJIS_Culiseta_flavivirus_strain_502-13 (NC_030290.1)
FJIS_Cell_fusing_agent_virus_strain_2 (MH237596.1)
FJIS_Cell_fusing_agent_virus (LR694076.1)
FJIS_Kamiti_River_virus (NC_005064.1)
FJIS_Aedes_flavivirus_strain_AEFV-SPFLD-MO-2011-MP6 (KC181923.1)
FJIS_Aedes_flavivirus_strain_Bangkok (KJ741266.1)
FJIS_Aedes_flavivirus_AEFVMQ29Bogor2017 (LC536088.1)
FJIS_Ochlerotatus_scapularis_flavivirus_bvmq18-25 (LC567152.1)
FJIS_Kilpisjarvi_flavivirus_isolate_FINL-201890 (ON949931.1)
FJIS_Xishuangbanna_aedes_flavivirus (NC_034017.1)
FJIS_Menghai_flavivirus_isolate_MHAedFV1 (NC_034204.1)
FJIS_Ochlerotatus_caspius_flavivirus_isolate_1608 (NC_034242)
FJIS_Hanko_virus (NC_030401)
FJIS_Parramatta_River_virus_isolate_92-B115745 (NC_027817.1)
FJIS_Mansonia_flavivirus_bvmq18-1 (LC567153.1)
FJIS_Tabanus_rufidens_flavivirus_18HF18 (LC540441.1)
FJUN_flavi_freshwater_ORF52_SRR12063566 (hou_et_al)
FJUN_XiangYun_flavivirus_isolate_XY61577 (OL700114.1)
FJAF_Photeros_flavivirus (OX394156.1)
FJAF_Sea_firefly_flavivirus (OX394161.1)
FJAF_Crangon_crangon_flavivirus (MK473878.1)
FJAF_Gammarus_chevreuxi_flavivirus_isolate_100_1 (MK473877.1)
FJAF_Gammarus_pulex_flavivirus_isolate_GPCM (MK473881.1)
FJAF_Harrimaniidae_flavivirus (OX394149.1)
FJAF_Cnidaria_flavivirus (OX394137.1)
FJFL_African_cichlid_flavi_like_virus (OX394152.1)
FJFL_Salmon_flavivirus_strain_Davis_UGA (MT075326.2)
FJFL_Tamana_bat_virus_isolate_P3_Vero (MZ229974.1)
FJFL_Cyclopterus_lumpus_virus_isolate_R170217-6L (MF776369.1)
FJFL_Southern_pygmy_squid_flavivirus_isolate_201405 (MK473875.1)
FJFL_Wenzhou_shark_flavivirus_strain_WZSS16343 (MG599986.1)
FJFL_Waxsystermes_virus (MW052131)
FJFL_Embiopteran_flavi-related_virus_isolate_OKIAV324 (MW208759.1)
FJFL_Chowder_bay_tunicate_associated_flavi_like_virus (OX394159.1)
FJFL_Flavivirus_sp._isolate_sequence_91 (MT949664.1)
FJUN_Iflavi_lake_water_ORF115_SRR6435398 (hou_et_al)
FJUN_flavi_salt_marsh_ORF170_SRR11829261 (hou_et_al)
FJUN_Fluviflavili_virus_isolate_PF1 (BK059737)
FJJI_Jingmen_tick_virus_isolate_WS3 (OM459837.1)
FJJI_Heilongjiang_tick_virus_isolate_HLJ41 (MK721860.1)
FJJI_Mogiana_tick_virus_isolate_Yunnan2016 (MT080097.1)
FJJI_Rhipicephalus_associated_flavi-like_virus_isolate_YNTV4 (MH814977.1)
FJJI_Jingmen_tick_virus_strain_JTVHAHH7 (MW721965.1)
FJJI_Jingmen_Tick_Virus_isolate_SY84 (NC_024113.1)
FJJI_Jingmen_tick_virus_isolate_JTMV_1 (MH155892.1)
FJJI_Jingmen_tick_virus_isolate_JTMV_3 (MH155896.1)
FJJI_Mogiana_tick_virus_isolate_MGTVV411 (NC_034222.1)
FJJI_Kindia_tick_virus_isolate_KITV20182 (MW341210)
FJJI_Jingmen_tick_virus_isolate_JMTVRh.microplusAm.variegatumFrench_Antilles (MN095523.1)
FJJI_Jingmen_tick_virus_isolate_TTP-Pool-3b (MN025512.1)
FJJI_Jingmen_tick_virus_isolate_TTP-Pool-19 (MN025516.1)
FJJI_Jingmen_tick_virus_isolate_JMTVPteropus_lyleiCambodia (MN095531.1)
FJJI_Takachi_virus_IM-OI110 (LC628196)
FJJI_Alongshan_virus_strain_Miass519 (MN648774.2)
FJJI_Guangdong_Jingmen-like_virus_isolate_118R-174486 (MW896893.1)
FJJI_Yanggou_tick_virus_strain_Erzin14-T20074 (MW525322)
FJJI_Hainan_jingmen-like_virus_isolate_s99-976258 (MW896920.1)
FJJI_Dipteran_jingmen-related_virus_isolate_OKIAV332 (MW314686.1)
FJJI_Trichopteran_jingmen-related_virus_isolate_OKIAV337 (MW314690.1)
FJJI_Wuhan_flea_virus_strain_WHZM (KR902713)
FJJI_Wuhan_aphid_virus_2_strain_WHYC-2 (KR902725)
FJJI_Wuhan_aphid_virus_1_strain_WHYC-1 (KR902721)
FJJI_Guaico_Culex_virus_isolate_LO35 (KM461666)
FJJI_Mole_Culex_virus_16GH38 (LC505052)
FJJI_Inopus_flavus_jingmenvirus_1_isolate_IFJV1 (OM869459)
FJJI_Wuhan_cricket_virus_strain_WHXS-1 (NC_028400)
FJUN_Infectious_precocity_virus_isolate_MR2018 (MT084113)
FJUN_flavi_wastewater_ORF139_SRR8857871 (hou_et_al)
0.2


Figure S2a

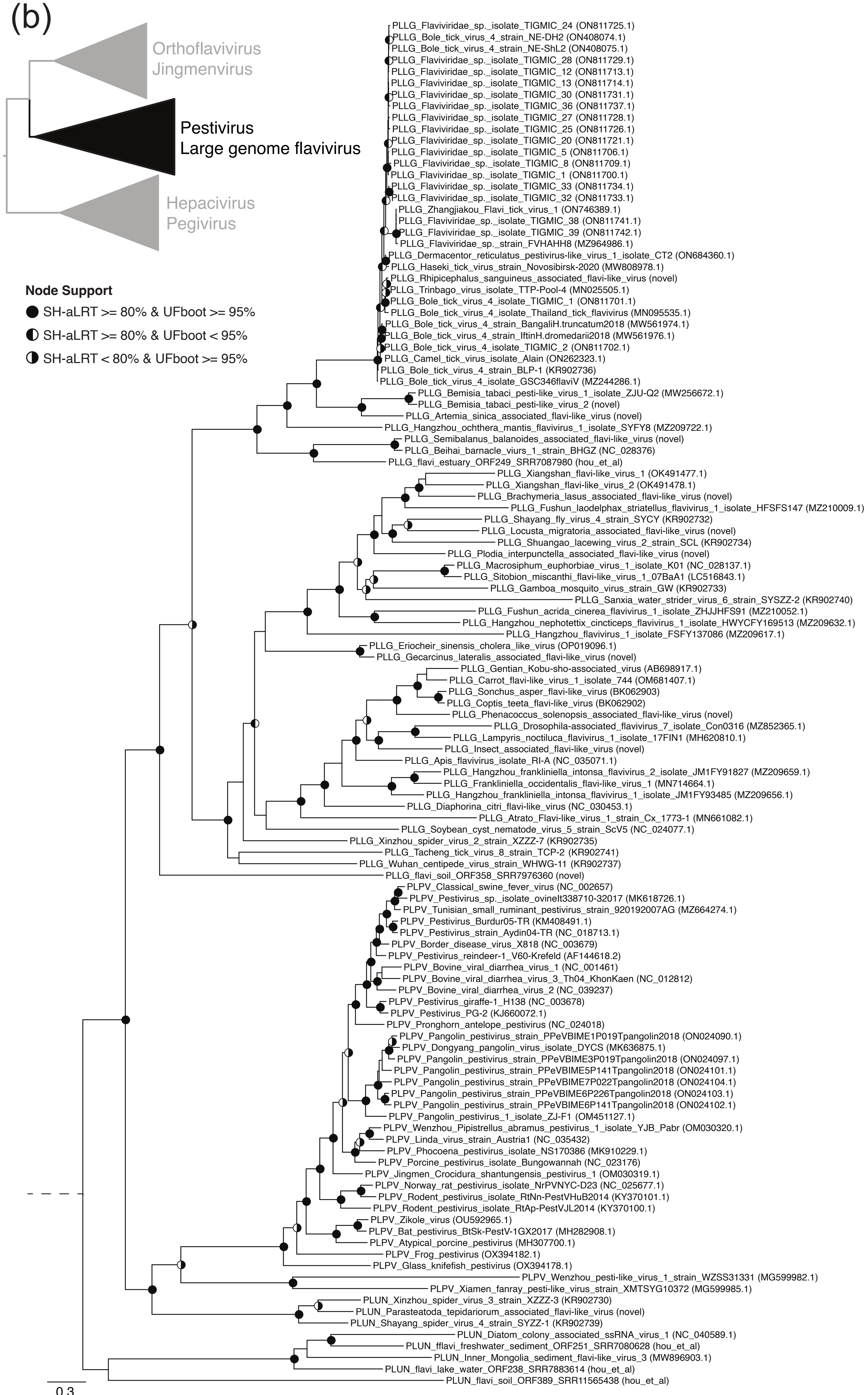
(b)
Orthoflavivirus
Jingmenvirus
Pestivirus
Large genome flavivirus
Hepacivirus
Pegivirus
Node Support
SH-aLRT >= 80% & UFboot >= 95%
SH-aLRT >= 80% & UFboot < 95%
SH-aLRT < 80% & UFboot >= 95%
PLLG_Flaviviridae_sp._isolate_TIGMIC_24 (ON811725.1)
PLLG_Bole_tick_virus_4_strain_NE-DH2 (ON408074.1)
PLLG_Bole_tick_virus_4_strain_NE-ShL2 (ON408075.1)
PLLG_Flaviviridae_sp._isolate_TIGMIC_28 (ON811729.1)
PLLG_Flaviviridae_sp._isolate_TIGMIC_12 (ON811713.1)
PLLG_Flaviviridae_sp._isolate_TIGMIC_13 (ON811714.1)
PLLG_Flaviviridae_sp._isolate_TIGMIC_30 (ON811731.1)
PLLG_Flaviviridae_sp._isolate_TIGMIC_36 (ON811737.1)
PLLG_Flaviviridae_sp._isolate_TIGMIC_27 (ON811728.1)
PLLG_Flaviviridae_sp._isolate_TIGMIC_25 (ON811726.1)
PLLG_Flaviviridae_sp._isolate_TIGMIC_20 (ON811721.1)
PLLG_Flaviviridae_sp._isolate_TIGMIC_5 (ON811706.1)
PLLG_Flaviviridae_sp._isolate_TIGMIC_8 (ON811709.1)
PLLG_Flaviviridae_sp._isolate_TIGMIC_1 (ON811700.1)
PLLG_Flaviviridae_sp._isolate_TIGMIC_33 (ON811734.1)
PLLG_Flaviviridae_sp._isolate_TIGMIC_32 (ON811733.1)
PLLG_Zhangjiakou_Flavi_tick_virus_1 (ON746389.1)
PLLG_Flaviviridae_sp._isolate_TIGMIC_38 (ON811741.1)
PLLG_Flaviviridae_sp._isolate_TIGMIC_39 (ON811742.1)
PLLG_Flaviviridae_sp._strain_FVHAHH8 (MZ964986.1)
PLLG_Dermacentor_reticulatus_pestivirus-like_virus_1_isolate_CT2 (ON684360.1)
PLLG_Haseki_tick_virus_strain_Novosibirsk-2020 (MW808978.1)
PLLG_Rhipicephalus_sanguineus_associated_flavi-like_virus (novel)
PLLG_Trinbago_virus_isolate_TTP-Pool-4 (MN025505.1)
PLLG_Bole_tick_virus_4_isolate_TIGMIC_1 (ON811701.1)
PLLG_Bole_tick_virus_4_isolate_Thailand_tick_flavivirus (MN095535.1)
PLLG_Bole_tick_virus_4_strain_BangaliH.truncatum2018 (MW561974.1)
PLLG_Bole_tick_virus_4_strain_IftinH.dromedarii2018 (MW561976.1)
PLLG_Bole_tick_virus_4_isolate_TIGMIC_2 (ON811702.1)
PLLG_Camel_tick_virus_isolate_Alain (ON262323.1)
PLLG_Bole_tick_virus_4_strain_BLP-1 (KR902736)
PLLG_Bole_tick_virus_4_isolate_GSC346flaviV (MZ244286.1)
PLLG_Bemisia_tabaci_pesti-like_virus_1_isolate_ZJU-Q2 (MW256672.1)
PLLG_Bemisia_tabaci_pesti-like_virus_2 (novel)
PLLG_Artemia_sinica_associated_flavi-like_virus (novel)
PLLG_Hangzhou_ochthera_mantis_flavivirus_1_isolate_SYFY8 (MZ209722.1)
PLLG_Semibalanus_balanoides_associated_flavi-like_virus (novel)
PLLG_Beihai_barnacle_viurs_1_strain_BHGZ (NC_028376)
PLLG_flavi_estuary_ORF249_SRR7087980 (hou_et_al)
PLLG_Xiangshan_flavi-like_virus_1 (OK491477.1)
PLLG_Xiangshan_flavi-like_virus_2 (OK491478.1)
PLLG_Brachymeria_lasus_associated_flavi-like_virus (novel)
PLLG_Fushun_laodelphax_striatellus_flavivirus_1_isolate_HFSFS147 (MZ210009.1)
PLLG_Shayang_fly_virus_4_strain_SYCY (KR902732)
PLLG_Locusta_migratoria_associated_flavi-like_virus (novel)
PLLG_Shuangao_lacewing_virus_2_strain_SCL (KR902734)
PLLG_Plodia_interpunctella_associated_flavi-like_virus (novel)
PLLG_Macrosiphum_euphorbiae_virus_1_isolate_K01 (NC_028137.1)
PLLG_Sitobion_miscanthi_flavi-like_virus_1_07BaA1 (LC516843.1)
PLLG_Gamboa_mosquito_virus_strain_GW (KR902733)
PLLG_Sanxia_water_strider_virus_6_strain_SYSZZ-2 (KR902740)
PLLG_Fushun_acrida_cinerea_flavivirus_1_isolate_ZHJJHFS91 (MZ210052.1)
PLLG_Hangzhou_nephotettix_cincticeps_flavivirus_1_isolate_HWYCFY169513 (MZ209632.1)
PLLG_Hangzhou_flavivirus_1_isolate_FSFY137086 (MZ209617.1)
PLLG_Eriocheir_sinensis_cholera_like_virus (OP019096.1)
PLLG_Gecarcinus_lateralis_associated_flavi-like_virus (novel)
PLLG_Gentian_Kobu-sho-associated_virus (AB698917.1)
PLLG_Carrot_flavi-like_virus_1_isolate_744 (OM681407.1)
PLLG_Sonchus_asper_flavi-like_virus (BK062903)
PLLG_Coptis_teeta_flavi-like_virus (BK062902)
PLLG_Phenacoccus_solenopsis_associated_flavi-like_virus (novel)
PLLG_Drosophila-associated_flavivirus_7_isolate_Con0316 (MZ852365.1)
PLLG_Lampyris_noctiluca_flavivirus_1_isolate_17FIN1 (MH620810.1)
PLLG_Insect_associated_flavi-like_virus (novel)
PLLG_Apis_flavivirus_isolate_RI-A (NC_035071.1)
PLLG_Hangzhou_frankliniella_intonsa_flavivirus_2_isolate_JM1FY91827 (MZ209659.1)
PLLG_Frankliniella_occidentalis_flavi-like_virus_1 (MN714664.1)
PLLG_Hangzhou_frankliniella_intonsa_flavivirus_1_isolate_JM1FY93485 (MZ209656.1)
PLLG_Diaphorina_citri_flavi-like_virus (NC_030453.1)
PLLG_Atrato_Flavi-like_virus_1_strain_Cx_1773-1 (MN661082.1)
PLLG_Soybean_cyst_nematode_virus_5_strain_ScV5 (NC_024077.1)
PLLG_Xinzhou_spider_virus_2_strain_XZZZ-7 (KR902735)
PLLG_Tacheng_tick_virus_8_strain_TCP-2 (KR902741)
PLLG_Wuhan_centipede_virus_strain_WHWG-11 (KR902737)
PLLG_flavi_soil_ORF358_SRR7976360 (novel)
PLPV_Classical_swine_fever_virus (NC_002657)
PLPV_Pestivirus_sp._isolate_ovineIt338710-32017 (MK618726.1)
PLPV_Tunisian_small_ruminant_pestivirus_strain_920192007AG (MZ664274.1)
PLPV_Pestivirus_Burdur05-TR (KM408491.1)
PLPV_Pestivirus_strain_Aydin04-TR (NC_018713.1)
PLPV_Border_disease_virus_X818 (NC_003679)
PLPV_Pestivirus_reindeer-1_V60-Krefeld (AF144618.2)
PLPV_Bovine_viral_diarrhea_virus_1 (NC_001461)
PLPV_Bovine_viral_diarrhea_virus_3_Th04_KhonKaen (NC_012812)
PLPV_Bovine_viral_diarrhea_virus_2 (NC_039237)
PLPV_Pestivirus_giraffe-1_H138 (NC_003678)
PLPV_Pestivirus_PG-2 (KJ660072.1)
PLPV_Pronghorn_antelope_pestivirus (NC_024018)
PLPV_Pangolin_pestivirus_strain_PPeVBIME1P019Tpangolin2018 (ON024090.1)
PLPV_Dongyang_pangolin_virus_isolate_DYCS (MK636875.1)
PLPV_Pangolin_pestivirus_strain_PPeVBIME3P019Tpangolin2018 (ON024097.1)
PLPV_Pangolin_pestivirus_strain_PPeVBIME5P141Tpangolin2018 (ON024101.1)
PLPV_Pangolin_pestivirus_strain_PPeVBIME7P022Tpangolin2018 (ON024104.1)
PLPV_Pangolin_pestivirus_strain_PPeVBIME6P226Tpangolin2018 (ON024103.1)
PLPV_Pangolin_pestivirus_strain_PPeVBIME6P141Tpangolin2018 (ON024102.1)
PLPV_Pangolin_pestivirus_1_isolate_ZJ-F1 (OM451127.1)
PLPV_Wenzhou_Pipistrellus_abramus_pestivirus_1_isolate_YJB_Pabr (OM030320.1)
PLPV_Linda_virus_strain_Austria1 (NC_035432)
PLPV_Phocoena_pestivirus_isolate_NS170386 (MK910229.1)
PLPV_Porcine_pestivirus_isolate_Bungowannah (NC_023176)
PLPV_Jingmen_Crocidura_shantungensis_pestivirus_1 (OM030319.1)
PLPV_Norway_rat_pestivirus_isolate_NrPVNYC-D23 (NC_025677.1)
PLPV_Rodent_pestivirus_isolate_RtNn-PestVHuB2014 (KY370101.1)
PLPV_Rodent_pestivirus_isolate_RtAp-PestVJL2014 (KY370100.1)
PLPV_Zikole_virus (OU592965.1)
PLPV_Bat_pestivirus_BtSk-PestV-1GX2017 (MH282908.1)
PLPV_Atypical_porcine_pestivirus (MH307700.1)
PLPV_Frog_pestivirus (OX394182.1)
PLPV_Glass_knifefish_pestivirus (OX394178.1)
PLPV_Wenzhou_pesti-like_virus_1_strain_WZSS31331 (MG599982.1)
PLPV_Xiamen_fanray_pesti-like_virus_strain_XMTSYG10372 (MG599985.1)
PLUN_Xinzhou_spider_virus_3_strain_XZZZ-3 (KR902730)
PLUN_Parasteatoda_tepidariorum_associated_flavi-like_virus (novel)
PLUN_Shayang_spider_virus_4_strain_SYZZ-1 (KR902739)
PLUN_Diatom_colony_associated_ssRNA_virus_1 (NC_040589.1)
PLUN_fflavi_freshwater_sediment_ORF251_SRR7080628 (hou_et_al)
PLUN_Inner_Mongolia_sediment_flavi-like_virus_3 (MW896903.1)
PLUN_flavi_lake_water_ORF238_SRR7883614 (hou_et_al)
PLUN_flavi_soil_ORF389_SRR11565438 (hou_et_al)
0.3

Figure S2b

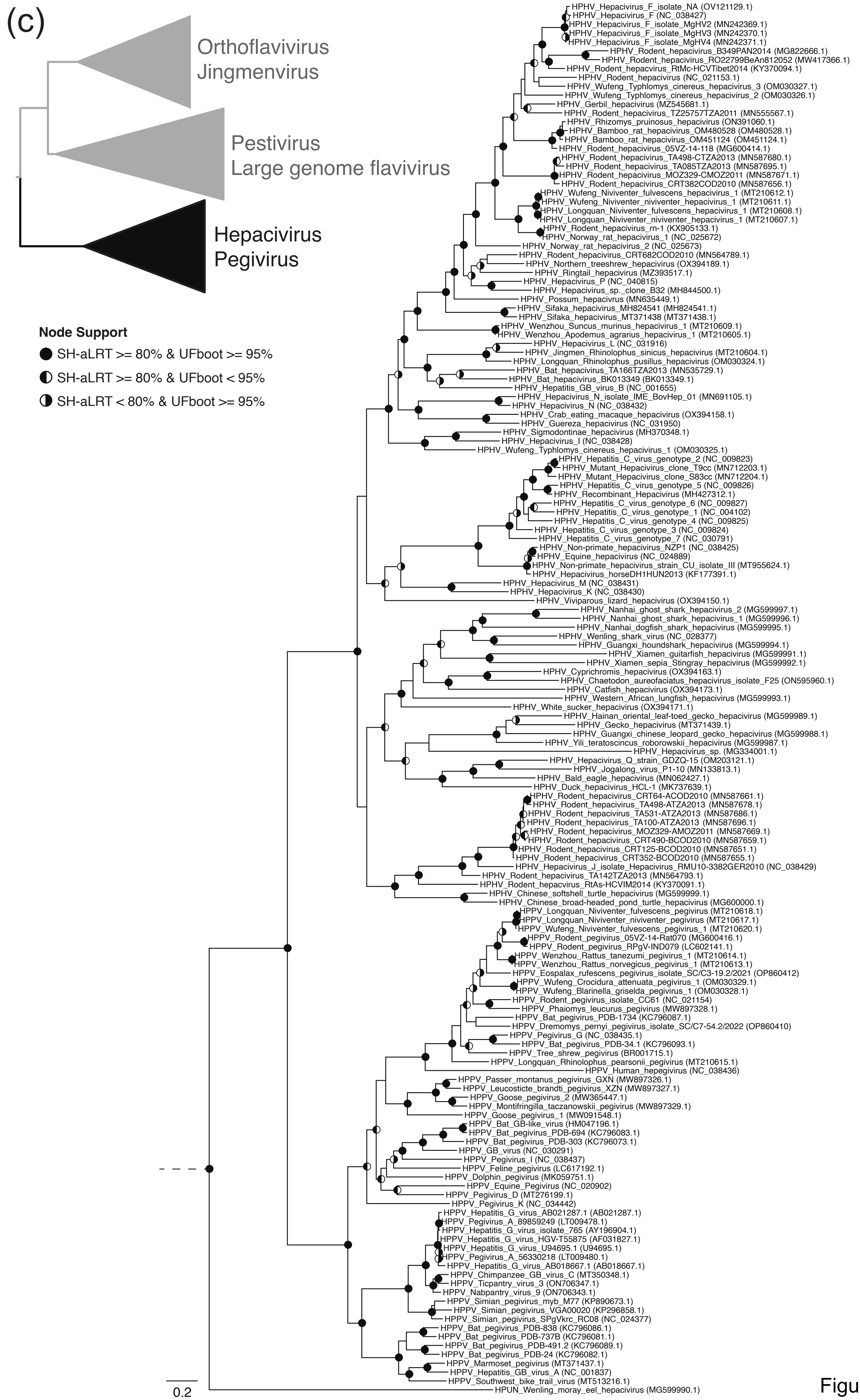
(c)
Orthoflavivirus
Jingmenvirus
Pestivirus
Large genome flavivirus
Hepacivirus
Pegivirus
Node Support
SH-aLRT >= 80% & UFboot >= 95%
SH-aLRT >= 80% & UFboot < 95%
SH-aLRT < 80% & UFboot >= 95%
HPHV_Hepacivirus_F_isolate_NA (OV121129.1)
HPHV_Hepacivirus_F (NC_038427)
HPHV_Hepacivirus_F_isolate_MgHV2 (MN242369.1)
HPHV_Hepacivirus_F_isolate_MgHV3 (MN242370.1)
HPHV_Hepacivirus_F_isolate_MgHV4 (MN242371.1)
HPHV_Rodent_hepacivirus_B349PAN2014 (MG822666.1)
HPHV_Rodent_hepacivirus_RO22799BeAn812052 (MW417366.1)
HPHV_Rodent_hepacvirus_RtMc-HCVTibet2014 (KY370094.1)
HPHV_Rodent_hepacivirus (NC_021153.1)
HPHV_Wufeng_Typhlomys_cinereus_hepacivirus_3 (OM030327.1)
HPHV_Wufeng_Typhlomys_cinereus_hepacivirus_2 (OM030326.1)
HPHV_Gerbil_hepacivirus (MZ545681.1)
HPHV_Rodent_hepacivirus_TZ25757TZA2011 (MN555567.1)
HPHV_Rhizomys_pruinosus_hepacivirus (ON391060.1)
HPHV_Bamboo_rat_hepacivirus_OM480528 (OM480528.1)
HPHV_Bamboo_rat_hepacivirus_OM451124 (OM451124.1)
HPHV_Rodent_hepacivirus_05VZ-14-118 (MG600414.1)
HPHV_Rodent_hepacivirus_TA498-CTZA2013 (MN587680.1)
HPHV_Rodent_hepacivirus_TA085TZA2013 (MN587695.1)
HPHV_Rodent_hepacivirus_MOZ329-CMOZ2011 (MN587671.1)
HPHV_Rodent_hepacivirus_CRT382COD2010 (MN587656.1)
HPHV_Wufeng_Niviventer_fulvescens_hepacivirus_1 (MT210612.1)
HPHV_Wufeng_Niviventer_niviventer_hepacivirus_1 (MT210611.1)
HPHV_Longquan_Niviventer_fulvescens_hepacivirus_1 (MT210608.1)
HPHV_Longquan_Niviventer_niviventer_hepacivirus_1 (MT210607.1)
HPHV_Rodent_hepacivirus_rn-1 (KX905133.1)
HPHV_Norway_rat_hepacivirus_1 (NC_025672)
HPHV_Norway_rat_hepacivirus_2 (NC_025673)
HPHV_Rodent_hepacivirus_CRT682COD2010 (MN564789.1)
HPHV_Northern_treeshrew_hepacivirus (OX394189.1)
HPHV_Ringtail_hepacivirus (MZ393517.1)
HPHV_Hepacivirus_P (NC_040815)
HPHV_Hepacivirus_sp._clone_B32 (MH844500.1)
HPHV_Possum_hepacivrus (MN635449.1)
HPHV_Sifaka_hepacivirus_MH824541 (MH824541.1)
HPHV_Sifaka_hepacivirus_MT371438 (MT371438.1)
HPHV_Wenzhou_Suncus_murinus_hepacivirus_1 (MT210609.1)
HPHV_Wenzhou_Apodemus_agrarius_hepacivirus_1 (MT210605.1)
HPHV_Hepacivirus_L (NC_031916)
HPHV_Jingmen_Rhinolophus_sinicus_hepacivirus (MT210604.1)
HPHV_Longquan_Rhinolophus_pusillus_hepacivirus (OM030324.1)
HPHV_Bat_hepacivirus_TA166TZA2013 (MN535729.1)
HPHV_Bat_hepacivirus_BK013349 (BK013349.1)
HPHV_Hepatitis_GB_virus_B (NC_001655)
HPHV_Hepacivirus_N_isolate_IME_BovHep_01 (MN691105.1)
HPHV_Hepacivirus_N (NC_038432)
HPHV_Crab_eating_macaque_hepacivirus (OX394158.1)
HPHV_Guereza_hepacivirus (NC_031950)
HPHV_Sigmodontinae_hepacivirus (MH370348.1)
HPHV_Hepacivirus_I (NC_038428)
HPHV_Wufeng_Typhlomys_cinereus_hepacivirus_1 (OM030325.1)
HPHV_Hepatitis_C_virus_genotype_2 (NC_009823)
HPHV_Mutant_Hepacivirus_clone_T9cc (MN712203.1)
HPHV_Mutant_Hepacivirus_clone_S83cc (MN712204.1)
HPHV_Hepatitis_C_virus_genotype_5 (NC_009826)
HPHV_Recombinant_Hepacivirus (MH427312.1)
HPHV_Hepatitis_C_virus_genotype_6 (NC_009827)
HPHV_Hepatitis_C_virus_genotype_1 (NC_004102)
HPHV_Hepatitis_C_virus_genotype_4 (NC_009825)
HPHV_Hepatitis_C_virus_genotype_3 (NC_009824)
HPHV_Hepatitis_C_virus_genotype_7 (NC_030791)
HPHV_Non-primate_hepacivirus_NZP1 (NC_038425)
HPHV_Equine_hepacivirus (NC_024889)
HPHV_Non-primate_hepacivirus_strain_CU_isolate_III (MT955624.1)
HPHV_Hepacivirus_horseDH1HUN2013 (KF177391.1)
HPHV_Hepacivirus_M (NC_038431)
HPHV_Hepacivirus_K (NC_038430)
HPHV_Viviparous_lizard_hepacivirus (OX394150.1)
HPHV_Nanhai_ghost_shark_hepacivirus_2 (MG599997.1)
HPHV_Nanhai_ghost_shark_hepacivirus_1 (MG599996.1)
HPHV_Nanhai_dogfish_shark_hepacivirus (MG599995.1)
HPHV_Wenling_shark_virus (NC_028377)
HPHV_Guangxi_houndshark_hepacivirus (MG599994.1)
HPHV_Xiamen_guitarfish_hepacivirus (MG599991.1)
HPHV_Xiamen_sepia_Stingray_hepacivirus (MG599992.1)
HPHV_Cyprichromis_hepacivirus (OX394163.1)
HPHV_Chaetodon_aureofaciatus_hepacivirus_isolate_F25 (ON595960.1)
HPHV_Catfish_hepacivirus (OX394173.1)
HPHV_Western_African_lungfish_hepacivirus (MG599993.1)
HPHV_White_sucker_hepacivirus (OX394171.1)
HPHV_Hainan_oriental_leaf-toed_gecko_hepacivirus (MG599989.1)
HPHV_Gecko_hepacivirus (MT371439.1)
HPHV_Guangxi_chinese_leopard_gecko_hepacivirus (MG599988.1)
HPHV_Yili_teratoscincus_roborowskii_hepacivirus (MG599987.1)
HPHV_Hepacivirus_sp. (MG334001.1)
HPHV_Hepacivirus_Q_strain_GDZQ-15 (OM203121.1)
HPHV_Jogalong_virus_P1-10 (MN133813.1)
HPHV_Bald_eagle_hepacivirus (MN062427.1)
HPHV_Duck_hepacivirus_HCL-1 (MK737639.1)
HPHV_Rodent_hepacivirus_CRT64-ACOD2010 (MN587661.1)
HPHV_Rodent_hepacivirus_TA498-ATZA2013 (MN587678.1)
HPHV_Rodent_hepacivirus_TA531-ATZA2013 (MN587686.1)
HPHV_Rodent_hepacivirus_TA100-ATZA2013 (MN587696.1)
HPHV_Rodent_hepacivirus_MOZ329-AMOZ2011 (MN587669.1)
HPHV_Rodent_hepacivirus_CRT490-BCOD2010 (MN587659.1)
HPHV_Rodent_hepacivirus_CRT125-BCOD2010 (MN587651.1)
HPHV_Rodent_hepacivirus_CRT352-BCOD2010 (MN587655.1)
HPHV_Hepacivirus_J_isolate_Hepacivirus_RMU10-3382GER2010 (NC_038429)
HPHV_Rodent_hepacivirus_TA142TZA2013 (MN564793.1)
HPHV_Rodent_hepacvirus_RtAs-HCVIM2014 (KY370091.1)
HPHV_Chinese_softshell_turtle_hepacivirus (MG599999.1)
HPHV_Chinese_broad-headed_pond_turtle_hepacivirus (MG600000.1)
HPPV_Longquan_Niviventer_fulvescens_pegivirus (MT210618.1)
HPPV_Longquan_Niviventer_niviventer_pegivirus (MT210617.1)
HPPV_Wufeng_Niviventer_fulvescens_pegivirus_1 (MT210620.1)
HPPV_Rodent_pegivirus_05VZ-14-Rat070 (MG600416.1)
HPPV_Rodent_pegivirus_RPgV-IND079 (LC602141.1)
HPPV_Wenzhou_Rattus_tanezumi_pegivirus_1 (MT210614.1)
HPPV_Wenzhou_Rattus_norvegicus_pegivirus_1 (MT210613.1)
HPPV_Eospalax_rufescens_pegivirus_isolate_SC/C3-19.2/2021 (OP860412)
HPPV_Wufeng_Crocidura_attenuata_pegivirus_1 (OM030329.1)
HPPV_Wufeng_Blarinella_griselda_pegivirus_1 (OM030328.1)
HPPV_Rodent_pegivirus_isolate_CC61 (NC_021154)
HPPV_Phaiomys_leucurus_pegivirus (MW897328.1)
HPPV_Bat_pegivirus_PDB-1734 (KC796087.1)
HPPV_Dremomys_pernyi_pegivirus_isolate_SC/C7-54.2/2022 (OP860410)
HPPV_Pegivirus_G (NC_038435.1)
HPPV_Bat_pegivirus_PDB-34.1 (KC796093.1)
HPPV_Tree_shrew_pegivirus (BR001715.1)
HPPV_Longquan_Rhinolophus_pearsonii_pegivirus (MT210615.1)
HPPV_Human_hepegivirus (NC_038436)
HPPV_Passer_montanus_pegivirus_GXN (MW897326.1)
HPPV_Leucosticte_brandti_pegivirus_XZN (MW897327.1)
HPPV_Goose_pegivirus_2 (MW365447.1)
HPPV_Montifringilla_taczanowskii_pegivirus (MW897329.1)
HPPV_Goose_pegivirus_1 (MW091548.1)
HPPV_Bat_GB-like_virus (HM047196.1)
HPPV_Bat_pegivirus_PDB-694 (KC796083.1)
HPPV_Bat_pegivirus_PDB-303 (KC796073.1)
HPPV_GB_virus (NC_030291)
HPPV_Pegivirus_I (NC_038437)
HPPV_Feline_pegivirus (LC617192.1)
HPPV_Dolphin_pegivirus (MK059751.1)
HPPV_Equine_Pegivirus (NC_020902)
HPPV_Pegivirus_D (MT276199.1)
HPPV_Pegivirus_K (NC_034442)
HPPV_Hepatitis_G_virus_AB021287.1 (AB021287.1)
HPPV_Pegivirus_A_89859249 (LT009478.1)
HPPV_Hepatitis_G_virus_isolate_765 (AY196904.1)
HPPV_Hepatitis_G_virus_HGV-T55875 (AF031827.1)
HPPV_Hepatitis_G_virus_U94695.1 (U94695.1)
HPPV_Pegivirus_A_56330218 (LT009480.1)
HPPV_Hepatitis_G_virus_AB018667.1 (AB018667.1)
HPPV_Chimpanzee_GB_virus_C (MT350348.1)
HPPV_Ticpantry_virus_3 (ON706347.1)
HPPV_Nabpantry_virus_9 (ON706343.1)
HPPV_Simian_pegivirus_myb_M77 (KP890673.1)
HPPV_Simian_pegivirus_VGA00020 (KP296858.1)
HPPV_Simian_pegivirus_SPgVkrc_RC08 (NC_024377)
HPPV_Bat_pegivirus_PDB-838 (KC796086.1)
HPPV_Bat_pegivirus_PDB-737B (KC796081.1)
HPPV_Bat_pegivirus_PDB-491.2 (KC796089.1)
HPPV_Bat_pegivirus_PDB-24 (KC796082.1)
HPPV_Marmoset_pegivirus (MT371437.1)
HPPV_Hepatitis_GB_virus_A (NC_001837)
HPPV_Southwest_bike_trail_virus (MT513216.1)
HPUN_Wenling_moray_eel_hepacivirus (MG599990.1)
0.2

Figure S2c

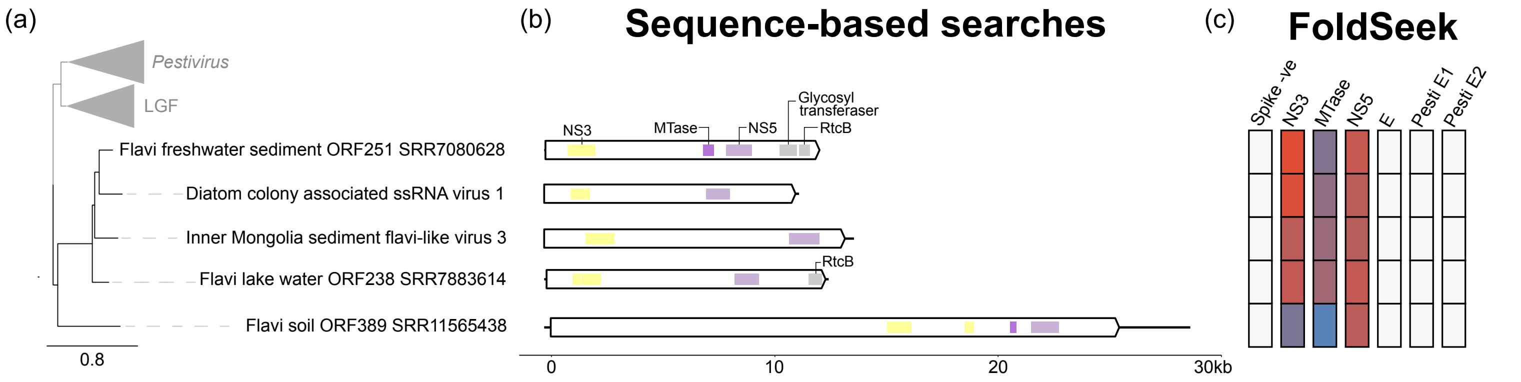
(a)
Pestivirus
LGF
Flavi freshwater sediment ORF251 SRR7080628
Diatom colony associated ssRNA virus 1
Inner Mongolia sediment flavi-like virus 3
Flavi lake water ORF238 SRR7883614
Flavi soil ORF389 SRR11565438
0.8
(b)
Sequence-based searches
NS3
MTase
NS5
Glycosyl transferaser
RtcB
RtcB
0
10
20
30kb
(c)
FoldSeek
Spike -ve
NS3
MTase
NS5
E
Pesti E1
Pesti E2

Figure S3
